# Angiotensin-(1-5) is a Potent Endogenous Angiotensin AT_2_-Receptor Agonist

**DOI:** 10.1101/2024.04.05.588367

**Authors:** Igor M. Souza-Silva, A. Augusto Peluso, Khalid Elsaafien, Antonina L Nazarova, Kasper B. Assersen, Lucas Rodrigues-Ribeiro, Mazher Mohammed, André F. Rodrigues, Arkadiusz Nawrocki, Lene Andrup Jakobsen, Pia Jensen, Annette D. de Kloet, Eric G. Krause, Mark Del Borgo, Ivan Maslov, Robert Widdop, Robson A. Santos, Michael Bader, Martin Larsen, Thiago Verano-Braga, Vsevolod Katritch, Colin Sumners, U. Muscha Steckelings

**Affiliations:** IMM - Department of Cardiovascular and Renal Research, University of Southern Denmark, Odense, Denmark; Neuroscience Institute, Georgia State University, Atlanta, GA, USA; Center for Neuroinflammation and Cardiometabolic Diseases, Georgia State University, Atlanta, USA; Department of Quantitative and Computational Biology, Center for New Technologies in Drug Discovery and Development, Bridge Institute, Michelson Center for Convergent Biosciences, University of Southern California, Los Angeles, CA, USA; Dept. of Dermatology, Odense University Hospital, Odense, Denmark; National Institute of Science and Technology in Nanobiopharmaceutics, Department of Physiology and Biophysics, Federal University of Minas Gerais (UFMG), Belo Horizonte, Brazil; Milton and Carroll Petrie Department of Urology, Icahn School of Medicine at Mount Sinai, New York, NY, USA; Max Delbrück Center for Molecular Medicine in the Helmholtz Association, Berlin, Germany; German Center for Cardiovascular Research (DZHK), Partner Site Berlin, Berlin, Germany; Department of Biochemistry and Molecular Biology, University of Southern Denmark, 5000 Odense, Denmark; Cardiovascular Disease Program, Biomedicine Discovery Institute, Department of Pharmacology, Monash University, Clayton, Victoria, Australia; Charité Universitätsmedizin Berlin, Corporate Member of Freie Universität Berlin and Humboldt-Universität zu Berlin, Berlin, Germany; University of Lübeck, Institute for Biology, Lübeck, Germany; Department of Physiology and Aging, College of Medicine, University of Florida, Gainesville, USA

**Author notes:** **Corresponding author:** Ulrike Muscha Steckelings, Institute for Molecular Medicine – Cardiovascular & Renal Research Unit University of Southern Denmark, Campusvej 55, 5230 Odense M, Denmark.

**Keywords:** angiotensin-(1-5), angiotensin AT_2_ receptor, phosphoproteomics, nitric oxide, blood pressure

## Abstract

**Background:** The renin-angiotensin system involves many more enzymes, receptors and biologically active peptides than originally thought. With this study, we investigated whether angiotensin-(1-5) [Ang-(1-5)], a 5-amino acid fragment of angiotensin II, has biological activity, and through which receptor it elicits effects.

**Methods:** The effect of Ang-(1-5) (1µM) on nitric oxide release was measured by DAF-FM staining in human aortic endothelial cells (HAEC), or Chinese Hamster Ovary (CHO) cells stably transfected with the angiotensin AT_2_-receptor (AT_2_R) or the receptor Mas. A potential vasodilatory effect of Ang-(1-5) was tested in mouse mesenteric and human renal arteries by wire myography; the effect on blood pressure was evaluated in normotensive C57BL/6 mice by Millar catheter. These experiments were performed in the presence or absence of a range of antagonists or inhibitors or in AT_2_R-knockout mice. Binding of Ang-(1-5) to the AT_2_R was confirmed and the preferred conformations determined by *in silico* docking simulations. The signaling network of Ang-(1-5) was mapped by quantitative phosphoproteomics.

**Results:** Key findings included: (1) Ang-(1-5) induced activation of eNOS by changes in phosphorylation at ^Ser1177^eNOS and ^Tyr657^eNOS and thereby (2) increased NO release from HAEC and AT_2_R-transfected CHO cells, but not from Mas-transfected or non-transfected CHO cells. (3) Ang-(1-5) induced relaxation of preconstricted mouse mesenteric and human renal arteries and (4) lowered blood pressure in normotensive mice – effects which were respectively absent in arteries from AT_2_R-KO or in PD123319-treated mice and which were more potent than effects of the established AT_2_R-agonist C21. (5) According to *in silico* modelling, Ang-(1-5) binds to the AT_2_R in two preferred conformations, one differing substantially from where the first five amino acids within angiotensin II bind to the AT_2_R. (6) Ang-(1-5) modifies signaling pathways in a protective RAS-typical way and with relevance for endothelial cell physiology and disease.

**Conclusions:** Ang-(1-5) is a potent, endogenous AT_2_R-agonist.

## INTRODUCTION

The renin-angiotensin system (RAS) is a complex hormonal system, in which the main effector hormone, angiotensin II (Ang II), is synthesized from its precursor angiotensinogen in a two-step enzymatic cascade catalyzed by renin and angiotensin converting enzyme (ACE) (1). This cascade was identified between the years 1898 and 1956. During the subsequent 30 years, the prevailing consensus was that with this discovery, the entire RAS had been identified.

However, since the late 1980’s and especially in recent years, many more angiotensin fragments have been characterized, which in some cases were found to be biologically active or even to have their own specific receptors. Among these fragments are angiotensin-(1-12) (2), angiotensin-(1-9) (3), angiotensin-(1-7) [Ang–(1-7)] (4,5), angiotensin III (6), angiotensin IV (7), and alamandine (8).

In contrast to the above listed fragments, the peptide angiotensin-(1-5) [Ang-(1-5)], which is cleaved from Ang-(1-7) by ACE, has been described to be a biologically inactive degradation product (9,10). However, its plasma levels are higher than those of Ang-(1-7) and have been shown to be regulated for example by addition of recombinant human ACE2 (rhACE2) [ACE2 cleaves Ang-(1-7) from Ang II] to human plasma samples (11) or in pathologies such as liver cirrhosis and COVID-19 (12,13).

These findings made us wonder whether such high amounts of Ang-(1-5) are really just “biological waste”, or whether this pentapeptide may have biological activity. Since Ang-(1-5) is derived from the main enzymatic cascade of the protective or alternative arm of the RAS (initiated by ACE2) (14), we hypothesized that Ang-(1-5) is a hormone of the protective RAS axis mediating actions such as nitric oxide (NO) release (15,16) and vasorelaxation (17).

For testing our hypothesis, we evaluated potential biological activity of Ang-(1-5) in various ways: measurement of NO release from human aortic endothelial cells (HAEC) *in vitro* , vascular effects using mouse mesenteric and human renal arteries *ex vivo* , and effects on blood pressure in normotensive C57BL/6 mice *in vivo* . Investigations aimed at identifying the receptor for Ang-(1-5) included studies on NO release in CHO-cells, which were transfected with either the angiotensin AT_2_-receptor (AT_2_R), or with the receptor Mas [the receptor for Ang-(1-7)]. Furthermore, interaction of Ang-(1-5) with the AT_2_R was modelled *in silico* by receptor docking modelling. The intracellular signaling pattern induced by Ang-(1-5) in HAEC was mapped by quantitative, time-resolved phosphoproteomics.

## METHODS

### Data Availability

The authors declare that the data supporting the findings of this study are available within the article and its Supplemental Material. The mass spectrometry proteomics data will be deposited to the ProteomeXchange Consortium via the PRIDE (18) partner repository and made publicly available when the manuscript will have been published in a peer reviewed journal.

Methodological details are provided in the Supplemental Material. Please see the Major Resources Table in the Supplemental Materials.

## RESULTS

### Proof of biological activity of angiotensin-(1-5)

#### Angiotensin-(1-5) activates eNOS and induces NO release from primary human aortic endothelial cells in vitro

As an initial approach to evaluate whether Ang-(1-5) is biologically active, we measured NO release from HAEC. Stimulation of HAEC with Ang-(1-5) (1µM) for 10 minutes led to a steady and statistically significant increase in DAF-FM fluorescence intensity indicating a continuous release of NO and biological activity of Ang-(1-5) (Fig. 1A).

**Figure 1.**
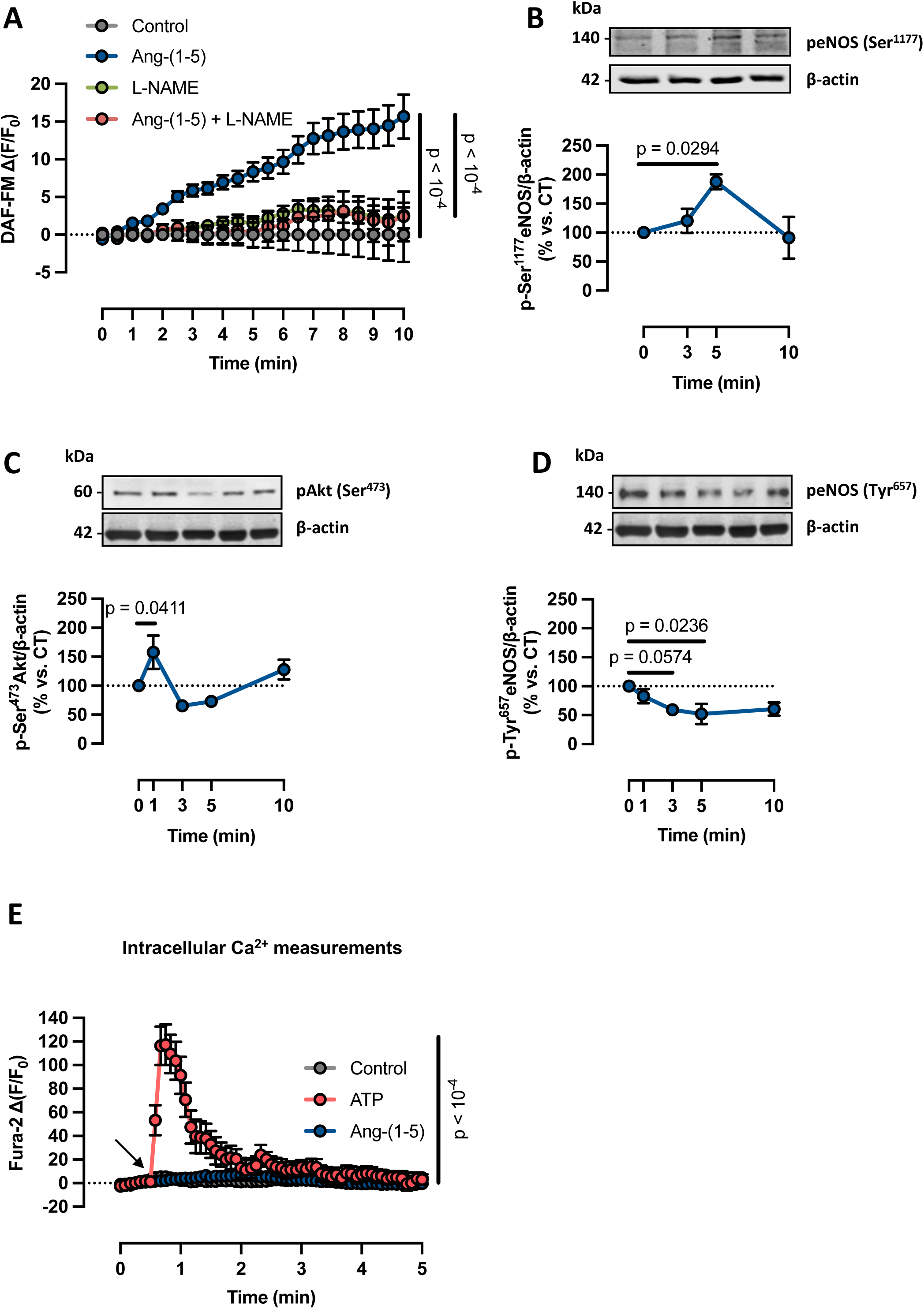
Effects of Ang-(1-5) on the regulation of NO release in vitro. **A**: Ang-(1-5) (1µM) significantly increased NO release in HAEC. The Ang-(1-5)-mediated increase in NO release was blocked by the NOS inhibitor L-NAME (10µM). **B**: Ang-(1-5) (1µM) significantly increased phosphorylation of eNOS at Ser^1177^ in HAEC after stimulation for 5 minutes. **C**: Phosphorylation of Akt at Ser^473^ in HAEC was significantly increased after 1 minute of stimulation with Ang-(1-5) (1µM). **D**: Ang-(1-5) (1µM) significantly reduced eNOS phosphorylation at Tyr^657^ after 5 minutes of stimulation with a strong trend already after 3 minutes. **E**: Ang-(1-5) (1µM) did not have any effect on intracellular Ca^2+^ levels in HAEC. Intracellular Ca^2+^ was significantly increased by the positive control, ATP (1µM). All data are shown as mean ± SEM from at least 3 independent experiments. Data on NO release and Ca^2+^ levels were analyzed by two-way repeated measures (RM)-ANOVA. Western Blot data on phosphorylation levels were analyzed by one-way ANOVA followed by Dunnett’s multiple comparison test. Differences were considered statistically significant when p ≤ 0.05. Exact p-values are provided within the figures.

NO is synthesized through cleavage of L-arginine by NO synthases (NOS). Therefore, we determined whether Ang-(1-5)-induced NO synthesis is indeed NOS dependent by testing the effect of Ang-(1-5) on NO release in the presence of the NOS inhibitor L-NAME (10µM). Since endothelial NOS (eNOS) is the main NOS isoform in endothelial cells, we further investigated eNOS-specific activation mechanisms such as phosphorylation of eNOS at serine^1177^ (Ser^1177^) by the kinase Akt or dephosphorylation of eNOS at tyrosine^657^ (Tyr^657^) (32).

Ang-(1-5)-induced NO synthesis was indeed completely blocked by L-NAME, thus suggesting that the increase in NO production by Ang-(1-5) was NOS dependent (Fig. 1A). Furthermore, we could show by Western Blotting that treatment of HAEC with Ang-(1-5) (1µM) induced eNOS activation mechanisms. Specifically, Ang-(1-5) led to a statistically significant increase in phosphorylation of Ser^1177^eNOS after 5 minutes of incubation (Fig. 1B). This was preceded by a statistically significant increase in Ser^473^-Akt phosphorylation after 1 minute of incubation with Ang-(1-5) (Fig. 1C), thus activating the kinase, which is responsible for Ser^1177^eNOS phosphorylation. Tyr^657^eNOS was significantly dephosphorylated after 5 minutes of incubation with a strong trend already present after 3 minutes (Fig. 1D). Ca^2+^-dependent pathways can alternatively activate eNOS (32), but are likely not involved in eNOS activation by Ang-(1-5), as Ang-(1-5) – in contrast to the positive control ATP (1 µM) - did not elicit statistically significant increases in intracellular Ca^2+^ in HAEC (Fig. 1E).

### Angiotensin-(1-5) does not mediate the effects of angiotensin-(1-7)

Since Ang-(1-5) is cleaved from Ang-(1-7) by angiotensin converting enzyme (ACE), we thought to test whether Ang-(1-7) has effects on its own at all or whether Ang-(1-5) is the true active component. For this purpose, we treated HAEC (which express the AT_2_R and Mas) with Ang-(1-7) (100nM) in the presence or absence of the ACE-inhibitor Captopril (100nM). Ang-(1-7) led to a significant increase in NO release from HAEC regardless of whether conversion of Ang-(1-7) to Ang-(1-5) was inhibited by Captopril (Suppl. Fig. S1) thus suggesting that effects of Ang-(1-7) are independent from generation of its metabolite Ang-(1-5).

#### Angiotensin-(1-5) induces relaxation of mouse and human resistance arteries ex vivo

Given that eNOS activation and the subsequent increase in NO release are classical promoters of vasodilation (33), we decided to evaluate whether Ang-(1-5) has vasorelaxant activity in resistance arteries *ex vivo* . Ang-(1-5) (1nM to 10µM) – under concomitant AT_1_R blockade by valsartan (3nM) – led to a significant and concentration-dependent relaxation of phenylephrine-preconstricted mouse mesenteric arteries (Fig.2A).

**Figure 2.**
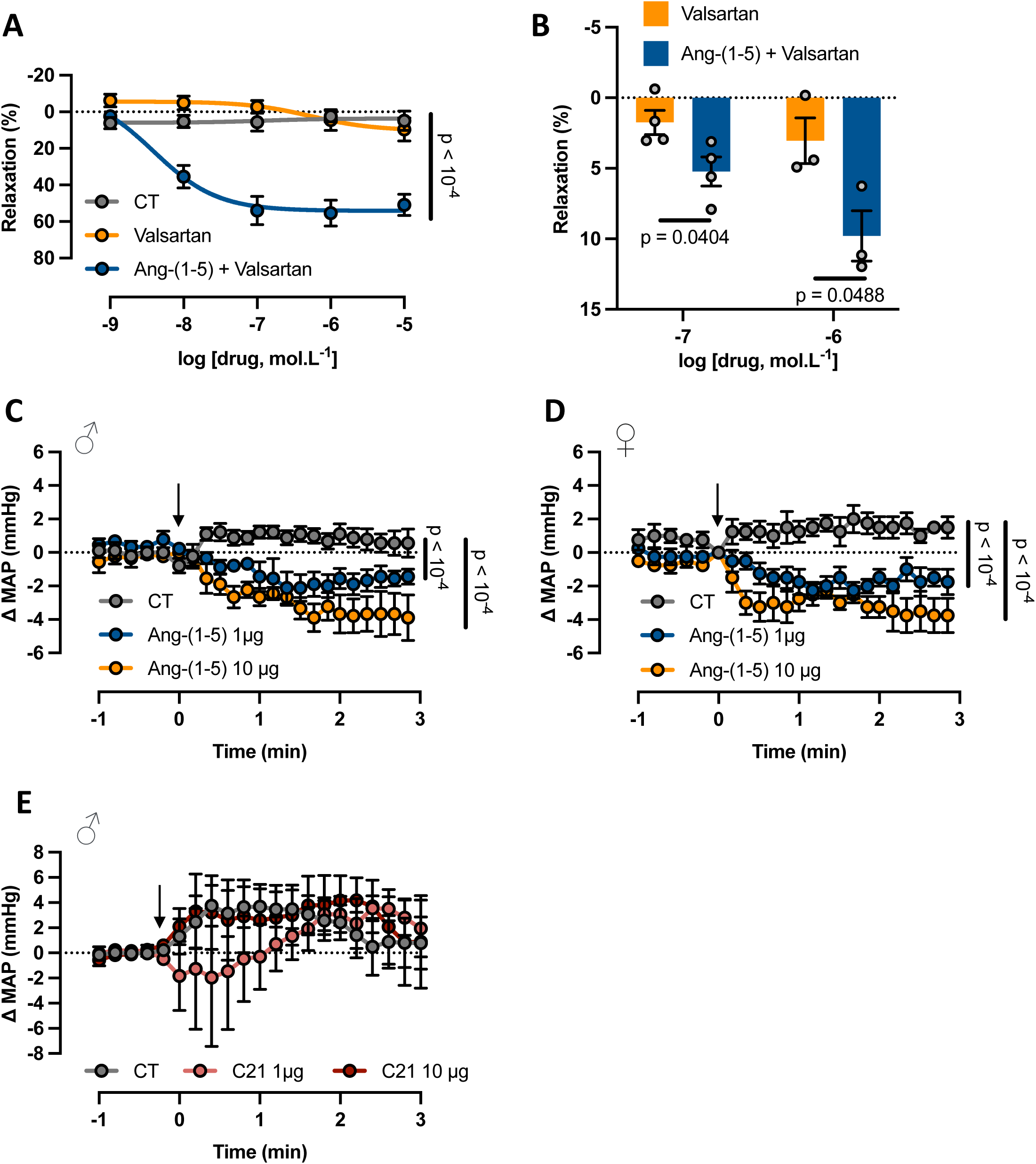
Vascular and BP effects of Ang-(1-5) **A**: Under AT_1_R blockade by valsartan (3nM), Ang-(1-5) induced a concentration-dependent, statistically significant relaxation of C57/Bl6 mouse mesenteric arteries pre-constricted with phenylephrine (1 µM). Valsartan (3nM) per se did not elicit changes in vasorelaxation. Segments that were pre-constricted with phenylephrine (1 µM) but not receiving any other treatment served as controls (CT). **B**: Under AT_1_R blockade by valsartan (3nM), Ang-(1-5) induced a significant relaxation of human renal arteries. **C,D**: A bolus injection of Ang-(1-5) (1 or 10µg iv) lowered blood pressure (BP) in anesthetized, normotensive (**C**) male and (**D**) female C57/Bl6 mice when compared to vehicle treated controls. **E**: C21 (1 or 10µg iv) did not have any BP-lowering effect in C57/Bl6 mice when compared to vehicle treated controls. Arrows indicate the timepoint of injections. All data are shown as mean ± SEM from at least 3 independent experiments (myography experiments) or from 9 male mice per each group (panel **C**), 4 female mice per each group (panel **D**) and 5 male mice per each group (panel **E**), respectively. Data on vasomotor effects in mouse mesenteric arteries were analyzed by two-way ANOVA. Data on vasomotor effects in human renal arteries were analyzed by multiple unpaired t-test. Data from blood pressure measurements were analyzed by two-way RM-ANOVA. Differences were considered statistically significant when p ≤ 0.05. Exact p-values are provided within the figures.

In order to test whether the vasorelaxant effect of Ang-(1-5) observed in mouse resistance arteries is also of relevance in humans, a potential vasorelaxant effect of Ang-(1-5) (under concomitant AT_1_R blockade by 3 nM valsartan to reliably prevent any AT_1_R-mediated vasoconstriction that could mask any AT_2_R effects) was evaluated in human renal arteries by wire myography. We indeed found that Ang-(1-5) (100nM or 1µM) led to a statistically significant, concentration-dependent relaxation of human renal arteries, thus strongly suggesting that Ang-(1-5) is also biologically active in humans (Fig. 2B).

#### Angiotensin-(1-5) lowers blood pressure in normotensive mice in vivo

Vasodilation of resistance arteries such as mesenteric arteries results in the lowering of blood pressure (BP) *in vivo* . Therefore, we evaluated, if bolus injections of Ang-(1-5) in normotensive, wild type C57BL/6J mice had any acute effects on BP. Ang-(1-5) (1µg or 10µg iv) led to a statistically significant, dose-dependent decrease in mean arterial pressure (MAP) in both male and female wild type C57BL/6J mice (Fig. 2 C,D). Of note, the established AT_2_R-agonist C21, applied iv at the same doses (1µg or 10µg), had no BP-lowering effect in male mice (Fig. 2E). Ang-(1-5) had no effect on heart rate (HR) in male C57BL/6J mice (Suppl. Fig. S2A). In female control mice, HR increased after iv injection of saline, which did not occur in Ang-(1-5)-treated mice and led to a statistically significant difference between the two groups (Suppl. Fig. S2B). C21 injections did not cause any change in HR (Suppl. Fig. S2C).

### Identification of the **AT_2_R** receptor as the receptor for angiotensin-(1-5)

Since Ang-(1-5) is an angiotensin fragment that shares its amino acid sequence with other RAS peptides such as Ang II or Ang-(1-7), we decided to evaluate if Ang-(1-5) may bind to and activate one of the known receptors of the RAS. We excluded the AT_1_R, since the effects of Ang-(1-5) we had seen so far, i.e. increased NO synthesis, vasodilation and lowering of BP are opposite to the known AT_1_R effects. To evaluate whether the AT_2_R or the receptor Mas mediates the effect of Ang-(1-5) on NO release, we used CHO cells stably transfected with either the AT_2_R (AT_2_R-CHO) or with the receptor Mas (Mas-CHO). Incubation with Ang-(1-5) (1µM) for 10 minutes induced NO release from AT_2_R-CHO cells, but not from Mas-CHO cells (Fig. 3A). To test functionality of the cells and accuracy of the assay, we used the AT_2_R agonist C21 (1µM) and the Mas agonist Ang-(1-7) (100nM) as positive controls. Both agonists increased NO release from cells with their respective target receptor [C21 in AT_2_R-CHO and Ang-(1-7) in Mas-CHO], but not from cells expressing the “non-target” receptor (Fig. 3A). Non-transfected CHO cells served as control to exclude any off-target effects of the compounds and were unresponsive to treatment with Ang-(1-5) (1µM), C21 (1µM) or Ang-(1-7) (100nM) (Fig. 3A). Collectively, these data strongly suggest that Ang-(1-5) is an agonist at the AT_2_R but not at the receptor Mas.

**Figure 3.**
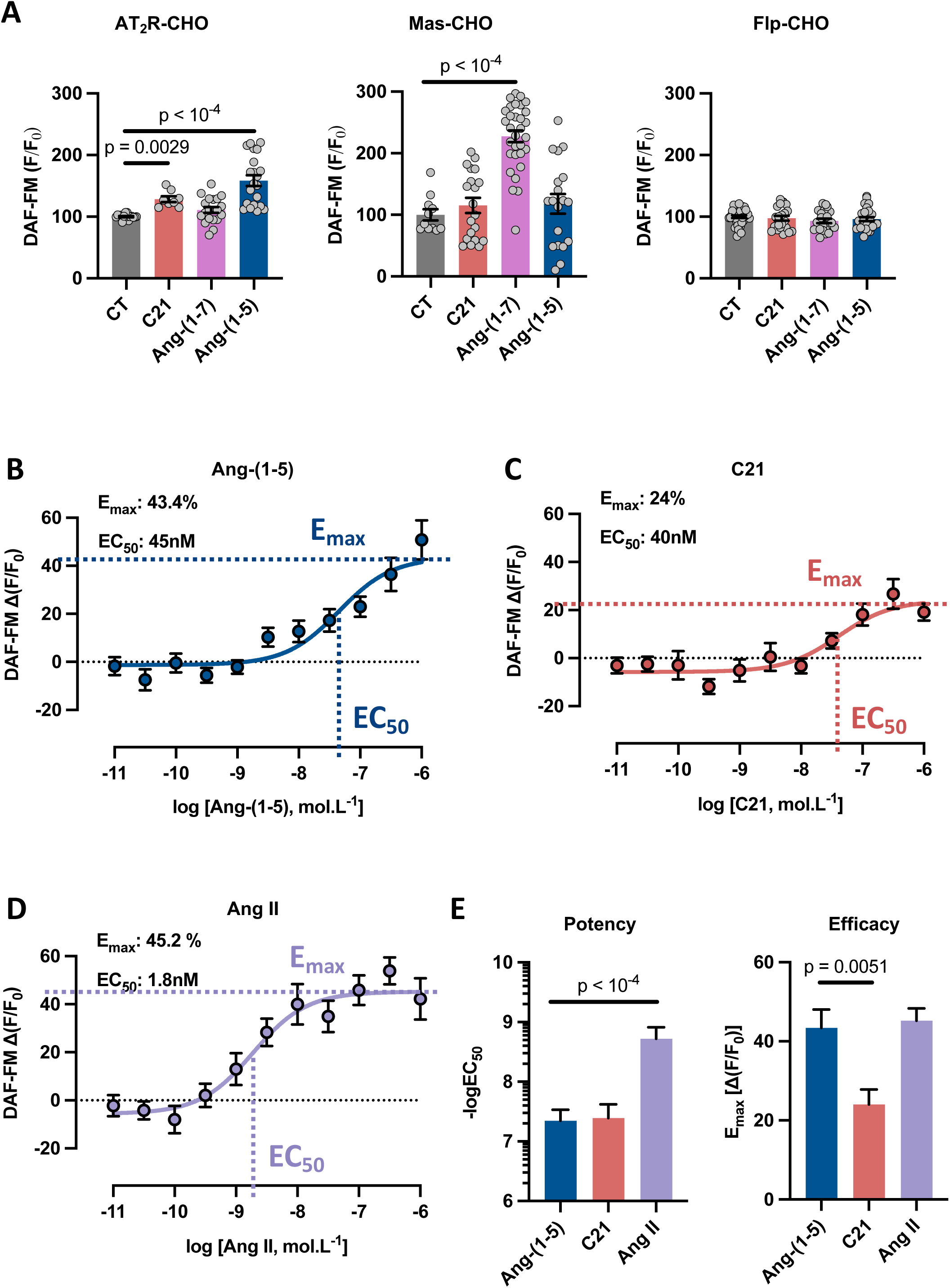
Evidence that Ang-(1-5) is a potent endogenous AT_2_R agonist **A:** Stimulation of AT_2_R-CHO with Ang-(1-5) (1µM) led to a statistically significant increase in NO release, as did stimulation with the AT_2_R agonist C21 (1µM, positive control). In Mas-CHO, stimulation with Ang-(1-5) (1µM) did not have any statistically significant effect on NO release. The Mas agonist Ang-(1-7) (100nM, positive control) significantly increased NO release. None of the compounds increased NO release in non-transfected CHO (Flp-CHO), suggesting that the responses observed were receptor-specific. Concentration-response curves for Ang-(1-5) (**B**), C21 (**C**) and Ang II (**D**) ranging from 10pM to 10µM were generated in AT_2_R-CHO revealing an EC_50_ of 45.97 ± 0.65nM and an E_max_ of 43.39 ± 4.64% for Ang-(1-5), an EC_50_ of 40.72 ± 0.59nM and an E_max_ of 24.06 ± 3.76% for C21 and an EC_50_ of 1.89 ± 0.65nM and an E_max_ of 45.21 ± 3.12% for Ang II. **E**: The EC_50_ for Ang-(1-5) was not statistically different from the EC_50_ for C21, but significantly lower than the EC_50_ for Ang II. The E_max_ for Ang-(1-5) was not statistically different from the E_max_ for Ang II and significantly higher than the E_max_ for C21. All data are shown as mean ± SEM from at least 3 independent experiments. Data were analyzed by one-way ANOVA followed by Dunnett’s multiple comparison test. Differences were considered statistically significant when p ≤ 0.05. Exact p-values are provided within the figures.

To further characterize the pharmacological properties of Ang-(1-5) at the AT_2_R, in particular efficacy (Emax) and potency (EC_50_), we generated concentration-response curves (CRCs) for the effect of Ang-(1-5) on NO release from AT_2_R-CHO cells. Incubation with Ang-(1-5) (10 pM to 10 µM) for 15 minutes led to a concentration-dependent increase in NO release with an E_max_ of 43.39 ± 4.64% and an EC_50_ of 45.97 ± 0.65 nM (Fig. 3B,E). To compare efficacy and potency of Ang-(1-5) with those of established AT_2_R agonists, we further constructed CRCs for C21 (10 pM to 10 µM) revealing an E_max_ of 24.06 ± 3.76% and an EC_50_ of 40.72 ± 0.59 nM (Fig. 3C,E) as well as for Ang II (10 pM to 10 µM) revealing an E_max_ of 45.21 ± 3.12% and an EC_50_ of 1.89 ± 0.65 nM (Fig. 3D,E). These data indicate that Ang-(1-5) has similar potency at the AT_2_R as C21 but lower potency than Ang II (Fig. 3E). Efficacy was comparable for Ang-(1-5) and Ang II, whereas the efficacy of Ang-(1-5) was significantly higher than that of C21 (Fig. 3E). Taken together, these results suggest that Ang-(1-5) is a potent AT_2_R agonist with high efficacy and pharmacological characteristics comparable to the established AT_2_R agonists Ang II and C21.

Further functional tests confirmed that Ang-(1-5) acts through the AT_2_R. Specifically, the vasorelaxant effect of Ang-(1-5) observed in segments from mesenteric arteries of C57BL/6 mice *ex vivo* (Fig. 2A,4A) was absent in arterial segments from AT_2_R deficient animals (Fig. 4A). *In vivo,* the BP-lowering effect of Ang-(1-5) (10 µg) observed in C57BL/6J mice (Fig. 2C,D) was abrogated by AT_2_R-blockade with PD123319 (100 µg) (Fig. 4B).

**Figure 4.**
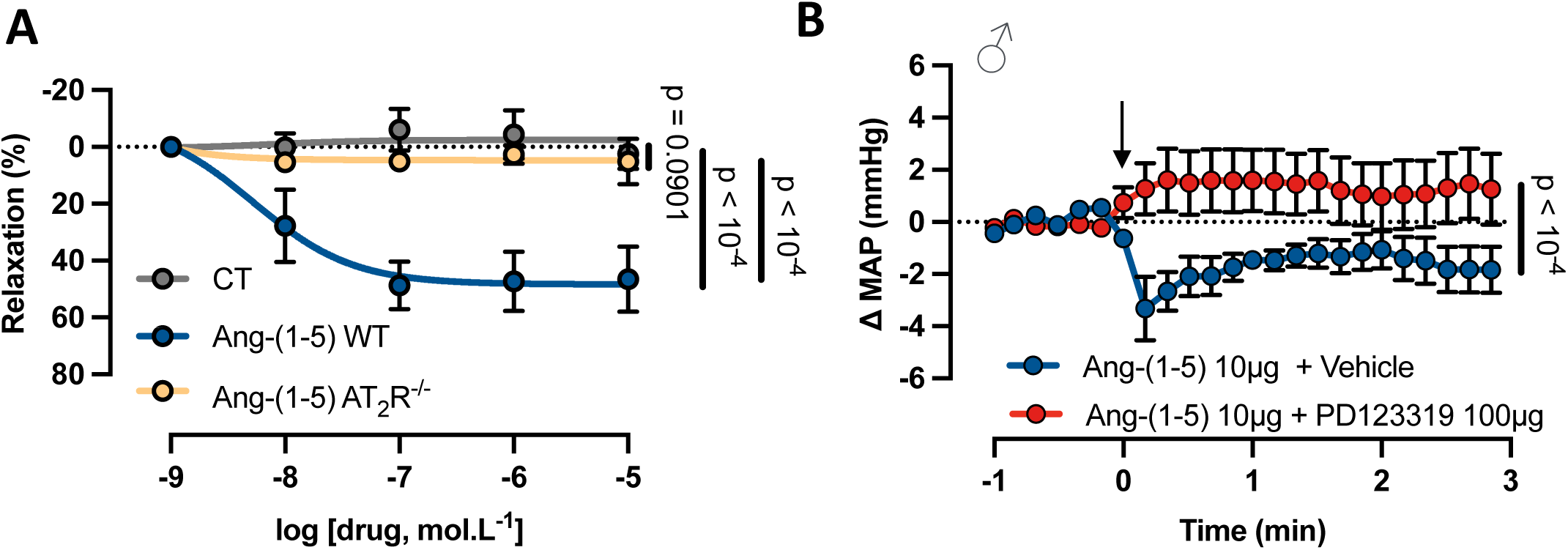
AT_2_R-dependence of Ang-(1-5) induced vasorelaxation and depressor effects. **A**: Relaxation of mouse mesenteric arteries pre-constricted with phenylephrine (1µM) in response to Ang-(1-5) (1nM to 10µM) was confirmed in arteries from C57/Bl6 wildtype (WT) mice but absent in arteries from AT R-knockout (AT_2_ R^-/-^) mice. All data are shown as mean ± SEM from at least 3 independent experiments. **B**: The BP-lowering effect by Ang-(1-5) (10µg) was blocked by the AT_2_R antagonist PD123319 (100µg). Arrow indicates the timepoint of Ang-(1-5) injection, with mice having been pretreated with 0.9% saline (vehicle) or PD123319 (100µg). All data are shown as mean ± SEM and are from 9 male mice (Ang-(1-5) + Vehicle group]), or 8 male mice (Ang-(1-5) + PD123319 group). Data were analyzed by two-way ANOVA (myography experiments) or two-way RM-ANOVA (BP measurements). Differences were considered statistically significant when p ≤ 0.05. Exact p-values are provided within the figures.

### Evidence for binding of angiotensin-(1-5) to the AT **_2_**R by *in silico* molecular docking simulations

To further support our previous findings, which suggested that Ang-(1-5) is an AT_2_R agonist, we evaluated if the peptide could interact with the AT_2_R by *in silico* molecular docking simulations. Molecular docking simulations revealed that Ang-(1-5) binds to the AT_2_R by two preferred, distinct conformations (Fig. 5A,B). In the first conformation (Conf1), Ang-(1-5) is superimposing the positioning of amino acids 1 to 5 of Ang II in the AT_2_R binding pocket (26) (Fig. 5A). In the second conformation (Conf2), Ang-(1-5) is located deeper in the AT_2_R binding pocket (Fig. 5B).

**Figure 5.**
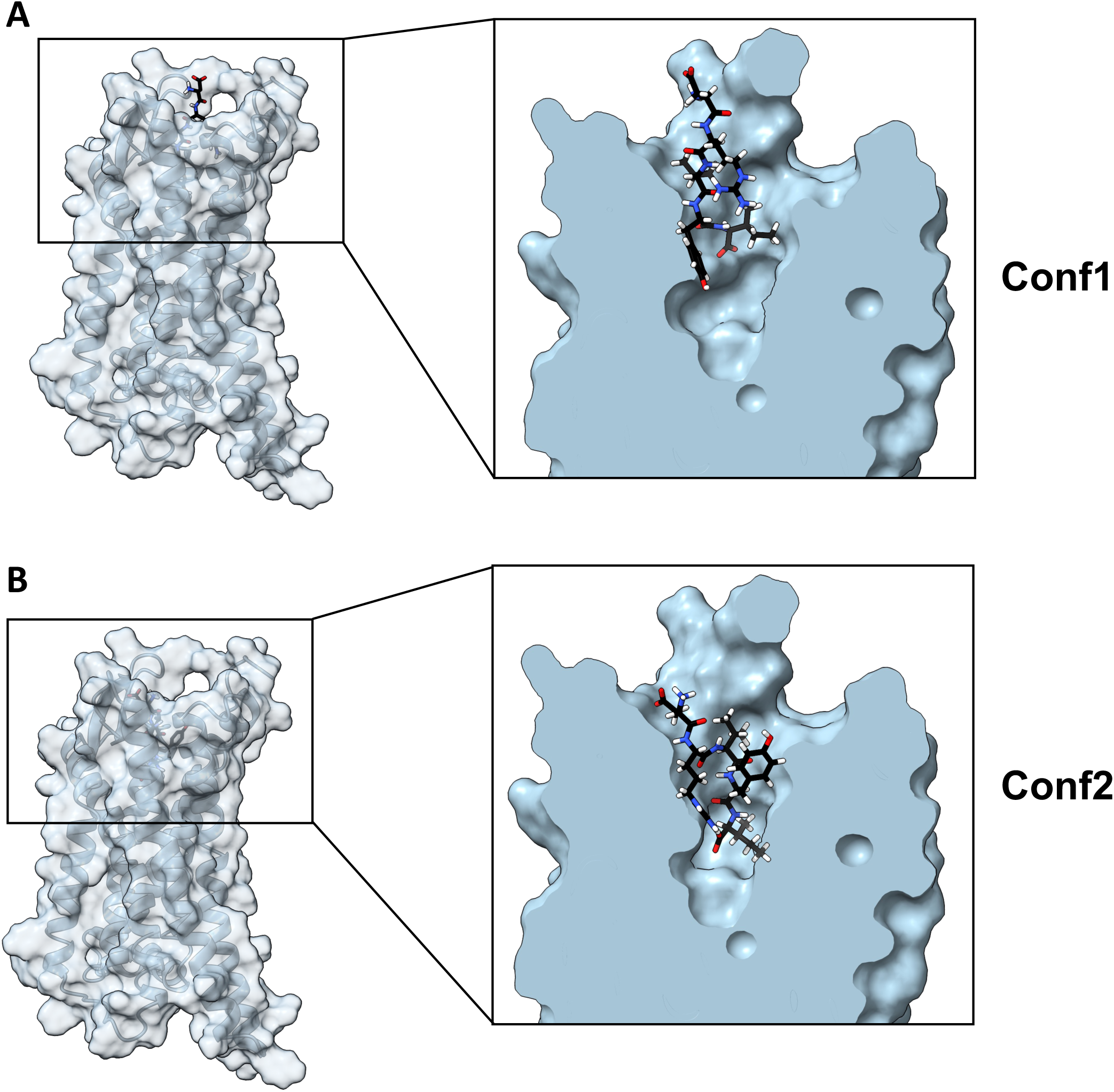
Molecular docking simulations of Ang-(1-5) binding to the AT_2_R. In silico simulation of docking of Ang-(1-5) to the structure of the activated AT_2_R revealed two preferred conformations for binding of Ang-(1-5) into the binding pocket. **A**: In Conformation 1, Ang-(1-5) is located and oriented in a similar way as Ang II in the binding pocket. **B**: In Conformation 2, Ang-(1-5) assumes a different orientation deeper in the binding pocket.

### Quantitative time-resolved phosphoproteomics reveals that angiotensin-(1-5) elicits signaling typical for the protective RAS

Mass spectrometry-based, time-resolved, quantitative phosphoproteomics was performed to explore the Ang-(1-5)-induced signaling network in HAEC based on the detection of changes in the phosphorylation status of the entire proteome. HAEC were stimulated with Ang-(1-5) (1µM) for 1, 3, 5 or 20 minutes. Resulting data were normally distributed (Suppl. Figs. 3,4). When compared to vehicle-treated controls, Ang-(1-5) led to a statistically significant modulation of phosphorylation levels of 1241 phosphopeptides derived by trypsin-digestion of 805 proteins. Such modifications occurred after treatment at all durations (1, 3, 5, and 20 minutes), with most changes after 20 minutes of stimulation (Fig. 6A,B). Ang-(1-5)-induced signaling in HAEC was mainly driven by dephosphorylations, which were more prevalent than phosphorylations at all timepoints tested (Fig 6B), a pattern that is typical for receptors and ligands of the protective RAS.

**Figure 6.**
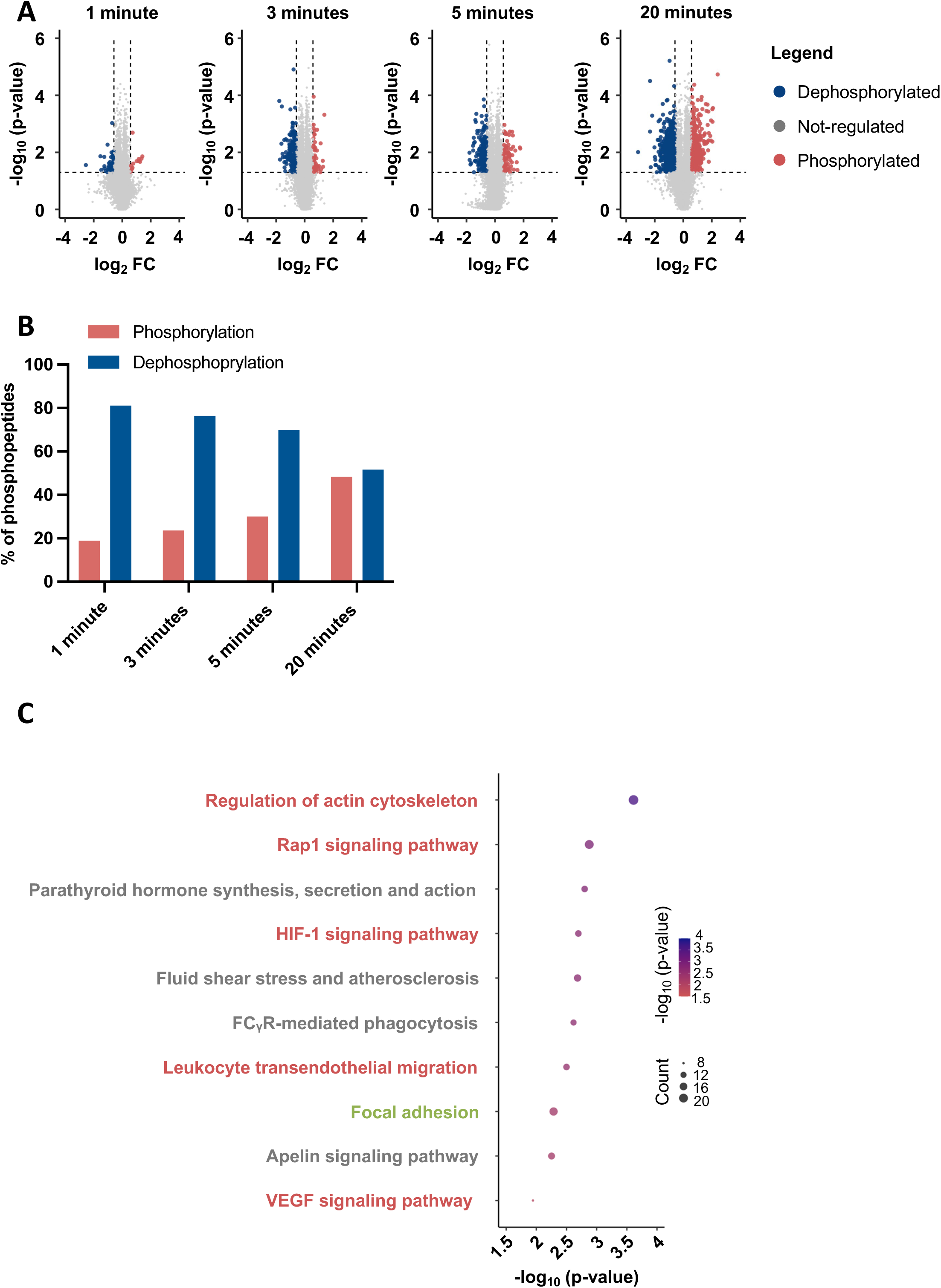
The profile of the Ang-(1-5) phosphoproteome in HAEC. **A**: Volcano-plots depicting log_10_ p-values and log_2_ fold-change (FC) of phospho-peptides identified by time-resolved, quantitative phosphoproteomics in HAEC treated with Ang-(1-5) for 1 minute, 3 minutes, 5 minutes or 20 minutes. Significance threshold was defined as p-value ≤ 0.05 and by a FC of ± 50%, depicted in the graphs by the dashed lines. Grey indicates non-regulated peptides. Blue indicates peptides in which phosphorylation levels were significantly decreased by Ang-(1-5) treatment. Red indicates peptides in which phosphorylation levels were significantly increased by Ang-(1-5) treatment. **B**: Percentage of phosphopeptide phosphorylations/dephosphorylations observed at every timepoint following treatment of HAEC with Ang-(1-5). **C**: KEGG pathway analysis of proteins with statistically significant modulations of phosphorylation levels by treatment with Ang-(1-5) in HAEC. Red indicates pathways inhibited by Ang-(1-5) treatment. Green indicates pathways activated by Ang-(1-5) treatment. Grey indicates pathways, for which Ang-(1-5)-mediated changes in phosphorylation levels could not be clearly assigned to pathway activation or inhibition. Statistical analysis of the enrichment of each term is expressed as -log_10_ p-value. Count represents the number of proteins comprised in each term.

Phosphoproteomics data were further analyzed by KEGG pathway analysis, which identifies signaling pathways that are enriched in response to a certain treatment, in our case Ang-(1-5). Figure 6C shows the top 10 signaling cascades enriched after KEGG analysis using only the significantly modified proteins in our phosphoproteomics data in response to Ang-(1-5) treatment. All of these signaling pathways are relevant for endothelial cell function and vascular disease. According to the phosphoproteomics data, pathways depicted in red are inhibited, pathways depicted in green are activated and for pathways depicted in grey the functional impact of Ang-(1-5) cannot be decided.

The potential impact of Ang-(1-5) on some of these pathways will be discussed in the Discussion. However, generally, phosphoproteomics for the identification of signaling mechanisms is a hypothesis-generating approach and any mechanisms suggested by bioinformatics analysis need to be verified by functional experiments. Nevertheless, the fact that all top 10 pathways are relevant for vascular physiology or disease supports our observations of a functional role of Ang-(1-5) in endothelial cells, the vasculature and BP control.

## DISCUSSION

Over the last four decades, it has become apparent that not only Ang II itself, but also many angiotensin fragments are biologically active, and that some even act through their own specific receptors. One fragment that has been regarded as biologically inactive is Ang-(1-5), a degradation product of Ang-(1-7) (9,10).

With this study, we are now providing evidence that Ang-(1-5) is a novel, biologically active hormone within the protective arm of the RAS. This evidence is based on the following key findings: (a) Ang-(1-5) induces NO release by activation of eNOS, (b) Ang-(1-5) relaxes preconstricted mouse and human resistance arteries, (c) Ang-(1-5) lowers BP in normotensive mice, (d) effects of Ang-(1-5) depend on the presence of AT_2_Rs and can be blocked with an AT_2_R antagonist, (e) according to molecular docking simulations, Ang-(1-5) binds to the AT_2_R in two preferred conformations, one of which is non-canonical, and (f) Ang-(1-5) induces a signal transduction pattern that is consistent with signaling networks identified for receptors of the protective RAS.

Ang-(1-5)-induced eNOS-dependent NO release resembles findings for other protective RAS compounds such as C21 (34), Ang-(1-7) (35) and alamandine (8), all of which were shown to activate eNOS resulting in NO synthesis. In all cases – including Ang-(1-5) – the eNOS activation mechanism involved phosphorylation of Ser^1177^, the canonical activation site, but not activation of the Ca^2+^/calmodulin pathway. As with C21, phosphatases are also likely involved in the Ang-(1-5) effect, as Ang-(1-5) dephosphorylated eNOS at Tyr^657^ – another activation mechanism (34).

Ang-(1-5)-induced NO release from endothelial cells *in vitro* translated into relaxation of mouse and human resistance arteries *ex vivo* . This result allowed for some additional conclusions, which were that (i) again, there was an effect typical for the protective RAS, (ii) Ang-(1-5) had a functional effect *ex vivo* , i.e. demonstrating biological activity in intact tissue. Together with the observation of *in vitro* NO release from **human** primary endothelial cells (HAEC), the Ang-(1-5)-induced relaxation of **human** kidney arteries strengthened evidence that effects of Ang-(1-5) are also relevant for the human situation. Evidence for relevance in humans is further supported by the presence of Ang-(1-5) in human plasma in amounts similar to other RAS hormones (36) and by the fact that Ang-(1-5) plasma levels undergo changes after pharmacological interference [AT1-receptor blockers (ARBs) (41), ACE2 (13)] or in disease (12).

We demonstrated *in vivo* relevance of Ang-(1-5) by showing a statistically significant Ang-(1-5)-induced decrease in BP in normotensive male and female C57Bl/6 mice. Although modest, this effect is in the same range as the effect of ARBs in normotensive animals (37). Remarkably, other protective RAS agonists such as C21 and Ang-(1-7) do not lower BP in normotensive animals as shown in this and other studies (24,38,39). Usually, BP-lowering effects of respective drugs including ARBs, C21 and Ang-(1-7) are stronger in hypertensive than in normotensive subjects (37). Whether this is also true for Ang-(1-5) will be investigated in future studies.

All of our experiments for the demonstration of biological activity of Ang-(1-5) *in vitro*, *ex vivo* and *in vivo* additionally aimed at identifying the receptor for Ang-(1-5). Since Ang-(1-5) induced NO release only in AT_2_R-CHO, but not in Mas-CHO, we concluded that the AT_2_R is likely the receptor for Ang-(1-5). This assumption was supported by the absence of Ang-(1-5)-induced vasorelaxation in mesenteric arteries from AT_2_R-deficient mice and by the inhibition of the BP lowering effect of Ang-(1-5) in mice by the AT_2_R-antagonist PD123319. Regarding NO release from AT_2_R-CHO, Ang-(1-5) displayed comparable potency (EC_50_) and higher efficacy (E_max_) than C21 and a 20-fold lower potency but a comparable efficacy as Ang II. Collectively, these data strongly suggest that Ang-(1-5) is a potent and selective AT_2_R agonist. The lack of effect of Ang-(1-5) in non-transfected CHO cells supported that the observed effects in AT_2_R-CHO were not off-target effects.

Demonstration of an interaction between the AT_2_R and Ang-(1-5) by *in silico* simulation of Ang-(1-5) docking to the crystal structure of the activated AT_2_R (26) further strengthened evidence that the AT_2_R is the receptor for Ang-(1-5). These simulations revealed two preferred conformations for Ang-(1-5) binding into the AT_2_R binding pocket. In conformation 1, Ang-(1-5) is superimposed with Ang II as described in the crystal structure obtained by Asada et al. (26). This conformation was somehow expected since Ang-(1-5) is a truncated analogue of Ang II. In the second conformation, Ang-(1-5) docked deeper into the AT_2_R binding pocket in a pose different from that described previously for any other AT_2_R ligand as part of an AT_2_R crystal structure (26,40–42). Further investigations are warranted to determine by which of these two poses Ang-(1-5) mainly elicits its biological effects and whether this is more dependent on a stable Ang-(1-5)/ AT_2_R interaction or on the interaction of Ang-(1-5) with structures deep in the receptor pocket. A unique, non-canonical mode of binding within the receptor pocket linked to a distinct activation mechanism may also be an explanation for the BP lowering effect of Ang-(1-5) in normotensive mice, which has not been observed in this and other studies for other AT_2_R agonists (24,38,39).

In order to explore signaling pathways elicited by Ang-(1-5) in HAEC, we applied mass spectrometry-based phosphoproteomics, which detects changes in the phosphorylation status of the entire cell proteome with many such changes being part of intracellular signaling cascades. Bioinformatic analysis revealed that the phosphorylation events modulated by Ang-(1-5) treatment were dephosphorylation-driven, which is unusual for conventional GPCRs such as the AT_1_R, but resembles signaling mechanisms identified by phosphoproteomics for other agonists/receptors of the protective RAS such as Ang-(1-7) (43), C21 (44) or alamandine (45). Analysis of the phosphoproteomics data by KEGG pathway analysis revealed that Ang-(1-5) seems to impact a number of signaling mechanisms with importance for endothelial cell and vascular physiology and pathophysiology. While proteomics data should be regarded as hypothesis-generating and need further investigation and verification, they still suggest for example an impact of Ang-(1-5) on vascular inflammation through effects on transendothelial migration and focal adhesion, an impact on endothelial cell migration and proliferation through effects on the cytoskeleton and VEGF signaling, or an impact on normal endothelial function and endothelial barrier integrity by interacting with Rap1 signaling. As mentioned before – all of these phosphoproteomics results are valuable hints towards Ang-(1-5) actions but warrant further investigations.

During the course of this study, another group published two articles reporting biological activity of Ang-(1-5) but claiming that Ang-(1-5) is a Mas agonist. The authors demonstrated an increase in atrial natriuretic peptide secretion from rat perfused atria and protective effects in a cardiac ischemia/reperfusion injury model (46,47), which confirms the protective nature of Ang-(1-5) as also observed by us. However, these investigations did not include a thorough characterization of Ang-(1-5) signaling and receptor interaction as our study. Furthermore, dosage of PD123319 was too low, and when concluding on the receptor for Ang-(1-5), the possibility of cross-inhibition due to Mas-AT_2_R heterodimerization was not considered (48). In our opinion, cross-inhibition makes interpretation of data using receptor antagonists or knockout animals more difficult than data from cells, which only express one of the two receptors as included in our study (e.g. AT_2_R-CHO or Mas-CHO). Nevertheless, while our data very strongly support that Ang-(1-5) is an AT_2_R agonist, the possibility that it also signals through Mas under certain conditions cannot be entirely excluded at this point and needs further investigation.

In summary, this study provides evidence that the angiotensin fragment Ang-(1-5) constitutes an endogenous AT_2_R agonist, which – when compared to Ang II – binds to the AT_2_R in a canonical and a non-canonical way. The signaling pattern induced by Ang-(1-5) is driven by dephosphorylations and is thus similar to signaling networks of other agonists of the protective RAS. The physiological actions of Ang-(1-5) identified so far are classical AT_2_R actions such as activation of eNOS resulting in increased NO synthesis in human endothelial cells, and relaxation of mouse mesenteric and human kidney arteries. In contrast to other AT_2_R agonists, Ang-(1-5) lowers BP in normotensive mice.

Thus, Ang-(1-5) can be regarded as a novel, potent AT_2_R agonist within the protective arm of the RAS with a role in cardiovascular physiology. Due to its non-canonical receptor binding, Ang-(1-5) may serve as a template for the design of a novel generation of drugs targeting the AT_2_R.

### Novelty and Significance (2-3 bullet points each)

#### What is known?

- The renin-angiotensin system harbors a variety of angiotensin peptides of different length, many, but not all of which are biologically active.
- Angiotensin-(1-5) has been largely regarded as a biologically inert degradation product of the active peptide angiotensin-(1-7).

#### What New Information Does the Article Contribute?

- Angiotensin-(1-5) is a biologically active, endogenous, potent agonist of the angiotensin AT_2_-receptor with an intracellular signaling pattern typical for ligands of the protective RAS.
- Angiotensin-(1-5) activates eNOS leading to an increase in nitric oxide release, to vasorelaxation and to a lowering of blood pressure in normotensive mice.
- Angiotensin-(1-5) binds to the AT_2_-receptor in a unique confirmation, which may give rise to effects different from those of other AT_2_-receptor agonists.

### Novelty and Significance

This study provides the first thorough characterization of Ang-(1-5) as a biologically active hormone of the RAS, specifically a potent AT_2_-receptor (AT_2_R) agonist. Biological effects identified for Ang-(1-5) in this study consist of activation of eNOS, synthesis of NO, relaxation of resistance arteries and lowering of blood pressure in normotensive mice. This range of effects clearly allocates Ang-(1-5) into the protective rather than the classical arm of the RAS. The signaling network elicited by Ang-(1-5) is typical for agonists of the protective RAS and bioinformatic analysis suggests effects on important mechanism of endothelial cell function and integrity such as migrations and proliferation, inflammation and cell adhesion or barrier function. *In silico* docking simulation confirmed binding of Ang-(1-5) to the AT_2_R and revealed that one of the preferred conformations within the binding pocket is fundamentally distinct from the conformation of Ang II in the binding pocket, which may constitute the basis for biased agonism at the AT_2_R. These different binding characteristics and the fact that Ang-(1-5) seems more potent than other AT_2_R agonists may make this molecule and its binding characteristics an interesting template for the design of new, more potent AT_2_R agonists.

## Nonstandard Abbreviations and Acronyms

ACE: Angiotensin Converting Enzyme
Akt: Protein Kinase Akt
Ang II: Angiotensin II
Ang-(1-5): Angiotensin-(1-5)
Ang-(1-7): Angiotensin-(1-7)
AT_2_R: Angiotensin AT_2_-receptor
AT_2_R: Angiotensin AT_1_-receptor
ATP: Adenosine Triphosphate
BP: Blood Pressure
C21: Compound 21
CHO: Chinese Hamster Ovary Cells
CRC: Concentration-Response Curve
EC_50_: Half-maximum effective concentration
E_max_: Maximum effect
HAEC: Human Aortic Endothelial Cells
HR: Heart Rate
iv: Intravenous
L-NAME: N^G^-Nitro-L-Arginine-Methyl Ester
MAP: Mean Arterial Pressure
NO: Nitric Oxide
NOS: Nitric Oxide Synthase
RAS: Renin-Angiotensin System
rhACE2: Recombinant human Angiotensin Converting Enzyme type 2
Ser: Serine
Tyr: Tyrosine

## ACKNOWLEDGMENTS

We thank the team of laboratory technicians at the Cardiovascular and Renal Research Unit, SDU, for their technical assistance.

## SUPPLEMENTAL MATERIAL

Supplemental Detailed Methods

Supplemental Figures S1 – S4

Major Resources Table

References 17 - 31 (included in reference list below)

