## Supplemental Methods for "Angiotensin-(1-5) is a Potent Endogenous Angiotensin AT_2_-Receptor Agonist"

**Supplemental, Detailed Methods**

**Cell culture**

Primary human aortic endothelial cells (HAEC) (Lonza, Switzerland) were grown in 25cm$^2$ culture flasks using Endothelial Cell Basal Medium-2 (EBM-2$^{TM}$) supplemented with EGM$^{TM}$-2 singleQuots® (Lonza, Switzerland). HAEC were used in passages 4 to 7. Chinese Hamster Ovary (CHO) cells carrying a Flp recombination target site (Invitrogen, USA) were stably transfected with the human $AT_2R$ ($AT_2R$-CHO), the human receptor Mas (Mas-CHO) or left non-transfected (Flp-CHO) (17). CHO cells were grown in 25cm$^2$ culture flasks using DMEM-F12 (Gibco, USA) supplemented with 10% fetal bovine serum (FBS; Gibco, USA) and 1% penicillin/streptomycin (Gibco, USA). G418 (100µg/mL) (Thermo Fisher Scientific, USA) was used for selection of $AT_2R$- or Mas-transfected cells. All cells were maintained at 37°C in a 5% $CO_2$ humified atmosphere.

**Nitric oxide release measurements**

***Nitric oxide release kinetics in HAEC***

Time-course experiments in HAEC were performed as previously described (19). Cells were cultured for 48 hours to 80% confluence on sterile, poly-L-lysine (Sigma, USA) coated glass coverslips of 10 mm diameter (0.13 – 0.16 mm thickness) (Thermo, USA), which were placed into 24-well plates (Nunc, USA). Cells were loaded for 30 minutes with the cell-permeable fluorescent NO indicator 4-amino-5-methylamino-2',7'-difluorofluorescein diacetate (1 µM; DAF-FM; excitation at 495nM, emission at 515nM) (Invitrogen, USA) diluted in serum-free and phenol red-free medium followed by washing with Hank's Balanced Salt Solution (HBSS). HAEC were stimulated with Ang-(1-5) (1µM) (Bachem, Switzerland), the NOS inhibitor $N_{\omega}$-nitro-L-arginine methyl ester (L-NAME) (10µM) (Sigma, USA) (20), or a combination of Ang-

(1-5) (1μM) and L-NAME (10μM). L-NAME was added to cells 10 minutes prior to addition of Ang-(1-5). Control groups were treated with equal amounts of vehicle (HBSS).

Fluorescence signals were recorded every 30 seconds over a period of 10 minutes by fluorescence microscopy (Olympus IX71, UK) with 20x zoom and FITIC laser exposure set to 200 ms. Full frame images were collected and analyzed for relative changes in fluorescence intensity (ΔF=F/F0) by *Xcellence* software (Olympus, UK). Only cells kept in focus during the whole experiment were analyzed. Correction for the weakening of the fluorescence signal caused by photobleaching, which is unrelated to the biological effect of interest, was done by normalizing data for each time point to the respective control values (same time point). To correct for differences in baseline signal intensity between independent experiments, control values for every time point were set to 100% and all other values presented as the difference from controls in percent. Statistical analysis was performed on the normalized original data.

***Nitric oxide release measurements in CHO cells***

For the identification of the receptor, which mediates the biological effects of Ang-(1-5), NO measurements were performed in $AT_2R$-CHO, Mas-CHO or Flp-CHO. Culturing of cells, DAF-FM loading, washing, microscopy, data acquisition and analysis followed the same protocol as mentioned above for HAEC except for the use of a different culture medium, namely DMEM-F12 supplemented with 10% FBS and 1% penicillin/streptomycin.

$AT_2R$-CHO or Mas-CHO, which do not express other RAS receptors, were treated with Ang-(1-5) (1μM) for 10 minutes to evaluate whether the $AT_2R$ or Mas mediates the effect of Ang-(1-5) on NO release as seen in HAEC. To ensure functionality of the transfected cells, $AT_2R$-CHO were stimulated with the $AT_2R$ agonist C21 (1μM) (kindly provided by Vicore Pharma, Sweden) and Mas-CHO with the Mas agonist Ang-(1-7) (100nM) (Bachem, Switzerland) for 10 minutes as positive controls. To ensure receptor specificity of agonist responses, $AT_2R$-CHO or Mas-CHO were further treated with the agonists for the respective other receptor [$AT_2R$-CHO with Ang-(1-7) (100nM) or Mas-CHO with C21 (1µM). Measurements in non-transfected Flp-CHO served to exclude any off-target effects of Ang-(1-5), C21 or Ang-(1-7). All agonists used were diluted in serum-free, phenol red-free HBSS. Cells treated with vehicle (serum-free and phenol red-free HBSS) served as controls.

***Generation of concentration-response curves in $AT_2R$-CHO***

To generate concentration-response curves, we performed a high-throughput, semi-automated NO release assay using $AT_2R$-CHO (20). 5000 cells per well were seeded onto 96 well plates and left to attach for 48 hours. Then, cells were loaded with DAF-FM diacetate (5µM) for 30 minutes in serum-free DMEM-F12. Concentration response curves for Ang-(1-5), C21 and Ang II (Sigma, USA) were generated, with concentrations ranging from 10pM to 1µM in 0.5 $\log_{10}$ steps. All drugs were diluted in serum-free DMEM-F12. Cells were stimulated for 15 minutes. Immediately after the stimulation period, cells were fixed with paraformaldehyde 4% (VWR, Denmark) for 5 minutes. Automated image acquisition was done by ImageXPress Pico fluorescence microscope device (Molecular Devices, USA) using a 4x magnification objective under excitation wavelength of 495nm and emission wavelength of 515nm. Automated image analysis was performed by the inbuilt ImageXPress Pico (Molecular Devices, USA) software. Data on fluorescence intensity from treated groups were normalized to the average of fluorescence intensity in control groups. Results are expressed in percentage increase from control.

For all protocols using NO release measurements as readout, at least 3 independent experiments were performed.

**Western Blotting**

***Treatment of cells***

Evaluation of eNOS phosphorylation was performed in subconfluent HAEC, which were serum-starved (0% FBS) for 4 hours prior to stimulation. Cells were stimulated with Ang-(1-5) (1µM) for 1, 3, 5 or 10 minutes. Vehicle-treated (serum-free HAEC medium) cells served as control.

***Isolation of proteins***

Cells were washed three times with ice-cold PBS and lysed by addition of 60µL lysis buffer [$Na_4P_2O_7$ 50mM, NaF 50mM, $Na_2$EDTA 5mM, NaCl 5mM, EGTA 5mM, HEPES 10mM, Triton X-100 0.5% and EDTA-free protease inhibitor cocktail (Roche, USA)] and mechanical scraping. The cell lysates were centrifuged at 14000 RPM for 20 min at 4$^{o}$C for cell debris removal. Protein concentration was quantified by Bradford Protein Assay Kit (BioRad, USA).

***Blotting, incubation, analysis***

A total of 12 µg protein was electrophoresed through a 10% bis-tris acrylamide gel and transferred to a nitrocellulose membrane (Thermo, USA). Membranes were blocked overnight in a solution containing 5% dry milk in Tris-buffered saline, 0.1% Tween-20 (Sigma, USA) (TBST) and incubated overnight at 4°C with the following primary antibodies: rabbit polyclonal anti-p-eNOS $Ser^{1177}$ (1:1000 – Cell Signaling, USA), rabbit monoclonal anti-p-eNOS $Tyr^{657}$ (1:1000 – Biotrend, UK), rabbit monoclonal anti-p-Akt $Ser^{473}$ (1:1000 – Cell Signaling, USA) or mouse monoclonal anti-β-actin (1:5000 – Abcam, USA). Membranes were incubated with secondary fluorescent IgG anti-rabbit (1:10.000, LI-COR, USA) or anti-mouse antibodies (1:10.000, LI-COR, USA) at room temperature for 1h, protected from light. Primary and secondary antibodies were diluted in TBST. Fluorescence signals were detected using the Odyssey infrared scanning system (LI-COR, USA) and quantified using Image Studio 4.0

software (LI-COR, USA). The intensity of fluorescence was normalized to β-actin levels and is presented as difference from controls in percent. For each time-point of stimulation, at least 3 independent experiments were performed.

**Measurement of intracellular $Ca^{2+}$ transients**

Subconfluent HAEC were cultivated on poly-L-lysine (Sigma, USA) coated glass coverslips prior to incubation with 1µM Fura-2-acetoxymethyl ester (Fura-2, excitation 340/380 nm, emission 510 nm, Thermo Scientific, USA) in serum-free medium for 30 minutes. Cells were washed and stabilized in 37°C HBSS. HAEC were treated with either Ang-(1-5) (1µM) or ATP (1µM) as positive control. Fluorescence signals were recorded every ten seconds over a period of 12 minutes by fluorescence microscopy (Olympus IX71) with 20x zoom and laser exposure set to 200 ms. The Fura-2 340/380 nm excitation fluorescence ratio was quantified using Xcellence software (Olympus, UK).

**Animals**

***Animals for myography***

Adult female C57BL/6J wild type mice were acquired from Charles River (Germany). Adult female $AT_2R$-knockout mice originated from the strain generated by Hein *et al*. (21). Mice were backcrossed repeatedly on a C57BL/6J background at the animal facilities of the University of Southern Denmark (Odense). Animals were housed in the animal facility at the Biomedical Laboratory in a controlled environment (12:12 hour light/dark cycle, 21 ± 3 °C, 55 ± 15 % humidity). Animals had free access to water and standard chow (#1324, Altromin, Denmark). Mice not bred at the facility were acclimatized for at least 5 days before they were included in a myograph experiment. Euthanasia of animals was approved by the Danish

Animal Experiments Inspectorate under the Danish Ministry of Justice, and all animal care followed the guidelines of the National Institutes of Health.

***Animals for cardiovascular measurements***

Studies were conducted on adult male and female C57BL/6J mice (Stock #000664, The Jackson Laboratory). Mice were 8-10 weeks old at the initiation of the studies. All animals were maintained in temperature and humidity-controlled rooms on 12:12 h light-dark cycles. Animals had free access to ad libitum food and water. All procedures were approved by the Institutional Animal Care and Use Committees at the University of Florida and were conducted in accordance with the National Institutes of Health's Guide for the care and use of laboratory animals and the ARRIVE guidelines (22). Surgery complications were predefined as an exclusion criterium. Such complications did not occur, and no animal had to be excluded. At the end of the experiment mice were euthanized by exposure to 5% isoflurane until 1 minute after breathing stops. Euthanasia was confirmed by decapitation.

**Wire myography**

***Myography in mouse mesenteric arteries***

*Preparation and mounting of arterial segments*

Mice were sacrificed by CO2 inhalation followed by severing of the carotid arteries. Intestines were isolated and instantly transported to the lab in physiological saline solution (PSS: NaCl 115mM, $NaHCO_3$ 25mM, $K_2HPO_4$ 2.5mM, $MgSO_4$ 1.2mM, HEPES 10mM, glucose 5.5mM, $CaCl_2$ 1.3mM) in a tube kept on ice. First-order mesenteric arteries from female WT C57BL/6J (10-14 weeks, n = 7) or female $AT_2R$-knockout mice on a C57BL/6J background (12-28 weeks, n = 7) were cleaned of perivascular adipose tissue and isolated with forceps and micro-scissors under a stereomicroscope. Segments of mesenteric arteries were then mounted in fourchamber wire myographs (40µm-diameter wires) from Danish Myo Technology (DMT, Denmark), submerged in PSS at 37°C, and constantly aerated with 5% CO2 in air. Time from death of the animal to completed mounting of the vessels was between 1 and 2 hours. Changes in isometric tension in arterial segments were recorded through an analog-to-digital converter (PowerLab 4/30 & Maclab 8e; ADInstruments, Australia) or directly from a myograph connected to a personal computer by the program Labchart Pro (ADInstruments). Prior to each experiment, the equipment was calibrated according to the manufacturer's instructions. Arterial segments rested for 30 minutes following the mounting procedure. Segments were normalized by establishing a length/tension relationship to determine the internal circumference corresponding to a passive transmural pressure of 100 mmHg ($IC_{100}$). Segments were then adjusted to an internal diameter corresponding to 90% of the internal circumference at IC100 which was used as IC1. Segments rested for another 30 minutes before proceeding.

*Arterial segment viability test*

Arterial segments were included in the experiment if they fulfilled the following criteria: 1. The segment contracted with a tension above 1N/m when challenged by an extracellular potassium concentration of 40 mM KPSS (PSS with 40 mmol/L KCl instead of 40 mmol/L NaCl), 2. The segment, when precontracted by 1 µM phenylephrine, relaxed more than 50 % in response to 10µM acetylcholine (where resting tension corresponds to 100% relaxation) and 3. The segment displayed stabile tension during precontraction with phenylephrine at a tension-level of < 50 % of 40mM KPSS induced contraction. Any segment which did not meet the required criteria was discarded.

*Test of mouse mesenteric artery vasorelaxation*

In order to investigate potential relaxing effects of Ang-(1-5), submaximal, amplitude-matched contractions (50% of the response to 32mM $K^+$) were induced using phenylephrine (1µM) as contractile stimulus (23). Experiments were performed under $AT_1R$ blockade by valsartan (3nM) (Sigma, USA) to reliably prevent any $AT_1R$-mediated vasoconstriction that could mask any $AT_2R$ effects (24). Valsartan was added 30 minutes prior to incubation with Ang-(1-5). Cumulative concentration-response curves were built with Ang-(1-5) (1nM to 10 µM). To confirm that any potential effect of Ang-(1-5) was mediated via the $AT_2R$, experiments were additionally performed in mesenteric artery segments from $AT_2R$ knockout mice and compared to responses in segments from wildtype mice. An additional arterial segment (time control), which was precontracted, but did not receive any other treatment, was used as control to determine spontaneous changes in contractile tension.

***Myography in human renal arteries***

In order to provide evidence that the observed effects of Ang-(1-5) are also relevant in humans, we performed myography experiments in human kidney arteries. The use of human kidney tissue was approved by the Ethical Committee of Southern Denmark (S-20100044); they were handled in agreement with the Declaration of Helsinki. Written informed consent was obtained from all patients before the experiments. All data collection was approved by the Danish Data Protection Agency.

*Isolation of human renal arteries*

Corticomedullary tissue for isolation of renal arcuate arteries was collected at the Department of Pathology. The tissue was isolated from human kidneys which were removed because of renal tumor growth. Special care was taken by the pathologist to isolate a tissue block from macroscopically healthy tissue. Tissue blocks were submerged in physiological saline solution on ice during transportation and dissection. Arcuate arteries were immediately isolated from the tissue block with forceps and microscissors under a microscope in a temperature-controlled environment (5°C). Isolated arcuate artery segments were stored in PSS in a closed container at 5 degrees Celsius over night for washout of anesthetics as previously described (25).

*Arterial segment viability test*

Arterial segments were included in the experiment if they fulfilled the following criteria: 1. The segment contracted with a tension above 1N/m when challenged by an extracellular potassium concentration of 40mM KPSS, 2. The segment, when precontracted by 40mM KPSS, relaxed more than 40% in response to 10µM acetylcholine (where resting tension corresponds to 100% relaxation). Any segment which did not meet the required criteria was discarded.

*Protocol for testing human renal artery vasorelaxation.*

In order to investigate potential relaxing effects of Ang-(1-5), submaximal, amplitude-matched contractions were induced using 15mM KPSS as contractile stimulus. Potential vasorelaxant effects of Ang-(1-5) were determined under concomitant $AT_1R$ blockade by valsartan (3nM), which was added 30 minutes prior to Ang-(1-5). Since submaximal contractions in human renal arteries were quite unstable and arteries tended to develop spontaneous oscillations after a few minutes, we did not run a cumulative concentration response curve as in mesenteric arteries, but only tested two concentrations of Ang-(1-5) (100nM and 1µM) – one at a time – in order to shorten the duration of the experiment. An additional arterial segment (time control), which was precontracted, but did not receive any other treatment, was used as control to determine spontaneous changes in contractile tension.

**Cardiovascular measurements in mice**

***Surgical procedures***

On the day of the experiment, anesthesia was induced in mice using isoflurane (3.5% in $O_2$) (USP, Dechra, USA) and maintained for the duration of the experiment with isoflurane (1.8% in $O_2$). Once animals were deeply anesthetized, a 1cm long segment of the right jugular vein was isolated from the surrounding tissue. Two silk sutures (size 5-0) were used to occlude blood flow from the caudal and rostral end. A puncture was made to the right jugular vein and a cannula (SAI Infusion Technologies, USA) was slid into the vein lumen and advanced to the right atrium. The catheter was then secured in place using the silk sutures. Following that, the left common carotid artery was isolated and separated from the vagal nerve. Silk sutures were used to occlude the artery and a puncture was made to advance a Millar catheter (Millar, USA) to the aortic arch. To acquire cardiovascular data, the Millar catheter was connected to a PowerLab signal transduction unit (ADInstruments, USA). Pulsatile arterial pressure was recorded at 1 kHz and blood pressure and heart rate calculated using Labchart8 software (ADInstruments, USA).

***Experimental Protocols***

Following the surgical procedures, cardiovascular parameters were allowed to stabilize for a 10-minute period followed by a 1-minute baseline recording. Pharmacological agents were then applied by intravenous bolus injection through the right jugular vein at a volume of 0.1ml and cardiovascular recordings performed for 3 minutes. Animals were assigned to treatment groups by the randomization function in Microsoft Excel. Investigators were blinded to the treatment.

*Angiotensin-(1-5)*

Angiotensin-(1-5) (Product #4030360, Bachem) was applied at 1 and 10µg to both male (n=9) and female (n=4) C57BL/6J mice. Each animal received both doses in a randomized order of administration. The first period of drug administration and recording was followed by a 30-minute washout period, after which the second intravenous administration was performed. Animals administered with vehicle (saline 0.9%; n=9 males and n=4 females) served as controls.

*Ang-(1-5) plus PD 123319*

$AT_2R$ involvement in BP effects elicited by Ang-(1-5) was evaluated by co-treatment with the $AT_2R$ antagonist PD123319 (Cat#1361, Tocris, USA) (100µg). For this experiment, male C57BL/6J mice were divided into two groups. In the first group, animals (n=9) were administered with a bolus injection of vehicle (saline 0.9%). Five minutes later, they were administered with a bolus injection of Ang-(1-5) (10µg) and recorded for three minutes. In the second group, animals (n=8) were administered with a bolus injection of PD123319 (100µg). Five minutes later, they were administered with Ang-(1-5) (10µg) and recorded for three minutes.

*Compound 21*

To compare the effect of Ang-(1-5) with that of an established $AT_2R$ agonist, Compound 21 (C21, Vicore Pharma, Sweden) was applied at 1 and 10µg to male C57BL/6J mice (n=5). As above, all doses were administered as bolus injection intravenously in the same animal, with the order of administration randomized between animals. Following administration, blood pressure was acquired for a period of 3 minutes, then a 30-minute washout period was applied before the second intravenous administration.

In all protocols, data was sampled at 12 second bins 1-minute before iv injections and 3-minutes following iv injections of drugs or control solution. For each animal, an average of the data collected during the 1-minute prior to iv injection was calculated as the baseline. At every 12 second bin, sampled data was then normalized to the baseline to calculate $\Delta$.

**Docking simulation of Ang-(1-5) binding to the $AT_2R$**

In order to provide additional evidence that Ang-(1-5) binds to the $AT_2R$, *in silico* docking simulations were performed. The $AT_2R$ protein structure bound to the endogenous agonist Ang II was derived from the RCSB server (https://www.rcsb.org; PDB: 6JOD) representing the active state of the receptor (26). The thermostability-improved apocytochrome $b_{562}$ (mbIIG) and Fab fragment of $AT_2R$-specific antibody (Fab4A03) were removed from the $AT_2R$ structure leaving the receptor protein subunit, and the crystallized Ang II peptide. The alanine $Ala^{208}$ was mutated back to serine $Ser^{208}$ and the protein was processed via the addition and optimization of hydrogens and optimization of the side chain residues.

Prior to conducting molecular docking, Ang-(1-5) underwent chiral definition and formal charge assignment. Ang-(1-5) molecular models were created from their two-dimensional representations, and their three-dimensional geometry was refined using the MMFF-94 force field (27). For docking simulations, a biased probability Monte Carlo (BPMC) optimization approach was employed, adjusting the internal coordinates of the compound based on pre-calculated grid energy potentials of the receptor (28). The grid potentials, while preserving the receptor conformational state, considered receptor flexibility through the usage of "soft" van der Waals potentials.

All-atom docking simulations were performed with the energy-minimized bond geometry of Ang-(1-5), and free torsion angles representing peptide flexibility, employing a sampling effort value of 30. The ligand docking box was selected to encompass the extracellular half of the protein for potential grid docking. At least 15 independent docking runs with 3 best conformations stored for each one were conducted, starting from random conformations. Consistency among the docking results was determined by comparing ligand conformations from the best ten docking poses. The unbiased docking procedure did not rely on distance restraints or any predefined information regarding the ligand-receptor interactions.

From these docking experiments, two top-scoring docking solutions, referred to as Conformation 1 and Conformation 2, representing Ang-(1-5) bound to $AT_2R$ complexes, were further refined. This refinement involved successive rounds of minimization and Monte Carlo sampling, focusing on the ligand conformation and including sidechain residues within 5 Å of the binding site. All the above-mentioned molecular modelling operations were performed in the ICM-Pro v3.9-2b molecular modelling and drug discovery suite (Molsoft LLC) (29).

**Mass spectrometry-based phosphoproteomics**

***Treatment and harvest of cells***

HAEC were stimulated with Ang-(1-5) (1μM) for 1, 3, 5, or 20 minutes. Cells treated with vehicle (serum-free HAEC medium) served as controls. Subsequently, cells were washed once with PBS followed by addition of 600μL of ice-cold phosphatase inhibitor solution (PhosSTOP, Roche Diagnostics, USA). Cells were removed by mechanical scraping and supernatant was transferred to 1.5mL Eppendorf tubes for centrifugation at 4°C, 300 x g for 5 minutes. Pellets were snap-frozen in liquid nitrogen and stored at -80°C until further usage.

***Protein isolation, reduction, alkylation and enzymatic digestion***

Cell pellets were resuspended in sodium deoxycholate (SDC) 1% in triethylammonium bicarbonate buffer (TEAB, pH 8). Cell pellets were tip-sonicated on ice twice for 10 seconds. 130μg of protein per sample was reduced with dithiothreitol (DTT) 10mM for 30 minutes. Proteins were alkylated by incubation with iodoacetamide (IAA) (20mM) for 20 minutes while protected from light. Enzymatic digestion of samples was performed by addition of endoproteinase Lys-C (0.01AU) (FUJIFILM Wako, USA) and trypsin (7.5μg; 1 part of trypsin to 50 parts of sample) (Promega, USA) to each tube followed by a 4-hour incubation at 37°C.

***TMT labelling***

TMTpro$^{TM}$ 16-plex (Thermo Fischer Scientific, USA) labelling reagent was reconstituted according to the manufacturer's instructions. Samples of tryptic peptides were labelled with the isobaric tags as follows: TMTpro126 for Control-1 (CT1), TMTpro127N for CT2, TMTpro127C for CT3, TMTpro128N for Ang-(1-5) treated cells for 1 minute-1 (1'1), TMTpro12C8N for 1'2, TMTpro129N for 1'3, TMTpro129C for Ang-(1-5) treated cells for 3 minutes-1 (3'1), TMTpro130N for 3'2, TMTpro130C for 3'3, TMTpro131N for Ang-(1-5) treated cells for 5 minutes-1 (5'1), TMTpro131C for 5'2, TMTpro132N for 5'3, TMTpro132C for Ang-(1-5) treated cells for 20 minutes-1 (20'1), TMTpro133N for 20'2, TMTpro133C for 20'3. TEAB (1M) was added to each tube to raise pH to around 8 followed by incubation of samples at room temperature for 90 minutes. Afterwards, samples were combined, and the resulting solution was lyophilized and stored at -80°C until further usage.

***Phosphopeptide enrichment***

Dried samples were reconstituted in loading buffer [acetonitrile 80%, trifluoroacetic acid 5% (TFA) and glycolic acid 1M]. To this solution, $TiO_2$ beads were added in a proportion of 0.6mg of beads per 100μg of proteins (starting material) and shaken vigorously for 10 minutes at room temperature. Subsequently, samples were centrifuged for 15 seconds at room temperature, the pellets were stored, and the supernatants were subjected to two more rounds of $TiO_2$ enrichment. At the end of this procedure, the supernatant (containing non-phosphorylated peptides) was lyophilized and stored at -80°C until further usage. $TiO_2$ beads from the three enrichment rounds were pooled and sequentially washed with buffer 1 (acetonitrile 80% and TFA 1%) and buffer 2 (acetonitrile 20% and TFA 0.1%). $TiO_2$ beads were incubated with elution buffer ($NH_4OH$ 1.25%, pH 11.3) and shaken for 10 minutes at room temperature. Samples were centrifuged, supernatants (containing phospho-enriched peptides) were collected and lyophilized until further usage. Non-phosphorylated and phospho-enriched peptides were reconstituted with TFA 0.1% and desalted using Oasis HBL Plus LP Extraction Cartridge (Waters, USA) or self-made Oligo™ R3 reversed-phase resin (Applied Biosystems, USA) columns, respectively. Desalted non-phosphorylated and phospho-enriched peptides were lyophilized and stored at -80°C until further usage.

***High-pH reverse-phase chromatography pre-fractionation***

Non-modified and phospho-enriched peptides were pre-fractionated by high-pH reverse phase chromatography. Briefly, non-phosphorylated and phospho-enriched peptides were reconstituted in buffer A [ammonium formate 20mM, pH 9.33) (Supelco, USA)] and loaded onto a Dionex Ultimate 300 (Thermo Fischer Scientific, USA) system. Chromatographic separation was performed in a nanoEase M/Z Peptide CSH C18 column (pore size 130Å, particle size 1.7µm, internal diameter 300µm, length 100mm) (Waters, USA), under a flow of 5µL/min and detection at 214nm. Mobile phase composition: buffer A ammonium formate 20mM, pH 9.33; buffer B ammonium formate 20mM, pH 9.33 20%, acetonitrile 80%. For non-phosphorylated peptides, 20 fractions were obtained with the following chromatographic gradient, expressed as % of buffer B: 0 to 14min 2%, 14 to 20min 2% to 10%, 20 to 85min 10% to 40%, 85 to 117min 40% to 50%, 117 to 122min 50% to 90%, 122 to 132min 90%, 132 to 133min 2%, 133 to 148min 2%. Fractions were collected from minute 15 every 151 seconds until minute 148. For phospho-enriched peptides, 12 fractions were obtained with the following chromatographic gradient, expressed as % of buffer B: 0 to 14min 2%, 14 to 74min 2% to 50%, 74 to 84min 50% to 70%, 84 to 89min 70% to 95%, 89 to 99min 95%, 99 to 100min

2%, 100 to 116min 2%. Fractions were collected from minute 15 every 151 seconds until minute 116. Fractions from non-phosphorylated and phospho-enriched peptides were lyophilized and stored at 80°C until further usage.

***Mass spectrometry analysis***

Non-phosphorylated and phospho-enriched peptides were reconstituted with 0.1% formic acid and loaded into an Easy-nLC 1000 liquid chromatographic system (Thermo Fischer Scientific, USA). Chromatographic separation was performed in a ReproSil-Pur 120 $C_{18}$-AQ column (pore size 120Å, particle size 1.9 µm, internal diameter 100 µm, length 200 mm) (Dr. Maisch GmbH, Germany), under a flow of 300nL/min. Mobile phase composition: buffer A 0.1% formic acid (Sigma, USA) in water; buffer B 95% acetonitrile, 0.1% formic acid. The chromatographic system was coupled to an Exploris 480 Orbitrap mass spectrometer (Thermo Fischer Scientific, USA), operating at positive mode with data-dependent acquisition. Non-phosphorylated and phospho-enriched peptides were separated with the following chromatographic method, expressed as % of buffer B: 0 to 100min 2 to 25%, 100 to 120min 25 to 40%, 120 to 121min 40% to 95%, 121 to 126min 95%, 126 to 127min 95% to 2%, 127 to 132min 2%. Eluting peptides were resolved at a resolution of 120000 FWHM at 200m/z with a mass range between 350 to 1200m/z. The top 10 most intense ions were selected for HCD fragmentation with isolation window of 0.7m/z and normalized collision energy of 33%. Resolution for MS2 was 45000 FMWH at 200m/z, with maximum injection time of 100ms for non-phosphorylated peptides and 200ms for phospho-enriched peptides.

The entire workflow of the phosphoproteomics experiment is illustrated below.

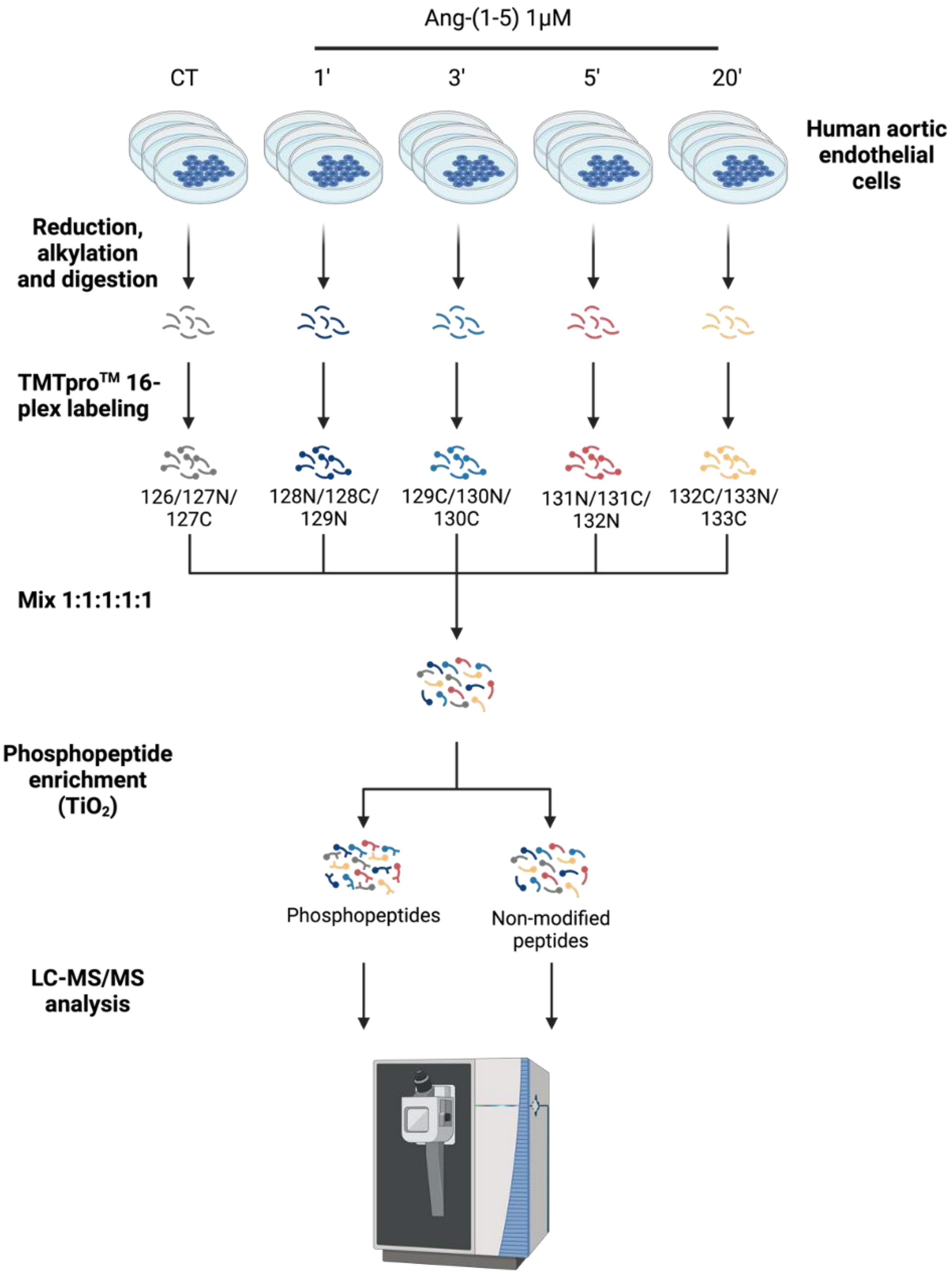


**Overview of phosphoproteomics experimental protocol**

HAEC were stimulated with Ang-(1-5) (1 μM) for 1, 3, 5 or 20 minutes or with vehicle (CT). Samples were lysed and proteins isolated for enzymatic digestion with trypsin and Lys-C. Digested peptides were labelled with TMTpro$^{TM}$ 16-plex and combined in a ratio of 1:1:1:1:1. Labelled and combined peptides underwent phosphopeptide enrichment with $TiO_2$. The resulting fractions containing phosphopeptides or non-modified peptides were analyzed by LC-MS/MS to generate the phosphoproteome and proteome datasets, respectively.

***Bioinformatic analysis***

Raw mass spectra (.raw files) were processed by Proteome Discoverer (Version 2.0, Thermo Fischer Scientific, USA) and submitted to MASCOT search engine against the Swiss-Prot *Homo sapiens* FASTA file. Trypsin and endoproteinase Lys-C were chosen as enzymes with allowance of up to two missed cleavages. In both non-phosphorylated and phospho-enriched sample analyses, carbamidomethyl (Cys) and TMTpro$^{TM}$ 16-plex (Lys and N-termini) were defined as fixed modifications. For non-phosphorylated and phospho-enriched samples, oxidation (Met) and deamidation (Asn) were defined as dynamic modifications. Additionally, in phospho-enriched samples, phosphorylation (Ser/Thr/Tyr) was also considered as a dynamic modification. Precursor mass tolerance of 5 ppm and fragment mass tolerance of 0.03 Da were applied. Quantification was performed only on peptides labelled with TMTpro$^{TM}$ 16-plex. Data were normalized by total peptide amount with a node in Proteome Discoverer. PhosphoRS node in Protein Discoverer was used to precisely identify phospho-site positions (30). Only peptides with a false discovery rate (FDR) below 1% and phosphopeptides with a site localization confidence ≥ 95% were considered for further analysis in this study. Data are expressed as $log_2$-transformed abundance ratio. Statistical analysis was performed by One-way ANOVA. Significance was considered when fold-change of $log_2$ abundance ratios was higher than ± 50% and p-value was ≤ 0.05.

Phospho-enriched $log_2$-transformed abundance ratios and *p*-values from each time-point were used to generate volcano-plots using the R package “ggplot2”. The number of significantly modulated phosphopeptides in response to Ang-(1-5) treatment was plotted in GraphPad Prism (v.10.0.0). KEGG (Kyoto Encyclopedia of Genes and Genomes) pathway database was used for protein pathway annotation and enrichment analysis in DAVID

Bioinformatics Resources 6.8 (https://david.ncifcrf.gov) (31) and results were plotted in R using the package “ggplot2”.

**Statistical analysis of data other than phosphoproteomics data**

All data were analyzed with GraphPad Prism 10.0. Data are presented as mean ± standard error of the mean (SEM). D’Agostino & Pearson normality test was applied to test data for normal distribution. Normally distributed data were analyzed with t-test, One-way ANOVA with Dunnett’s multiple comparison test, Two-way ANOVA or RM-Two-way ANOVA for comparison between independent groups. Repeated measures-ANOVA was used to analyze NO release kinetics and $Ca^{2+}$ transients in HAEC and measurements of blood pressure and heart rate in mice. For non-normally distributed data, Mann-Whitney test or Kruskal-Wallis test with Dunn’s multiple comparison test was performed. Statistical significance was considered when $p \leq 0.05$.
