## Supplemental Figures for "Angiotensin-(1-5) is a Potent Endogenous Angiotensin AT_2_-Receptor Agonist"

**Supplement – Figures**

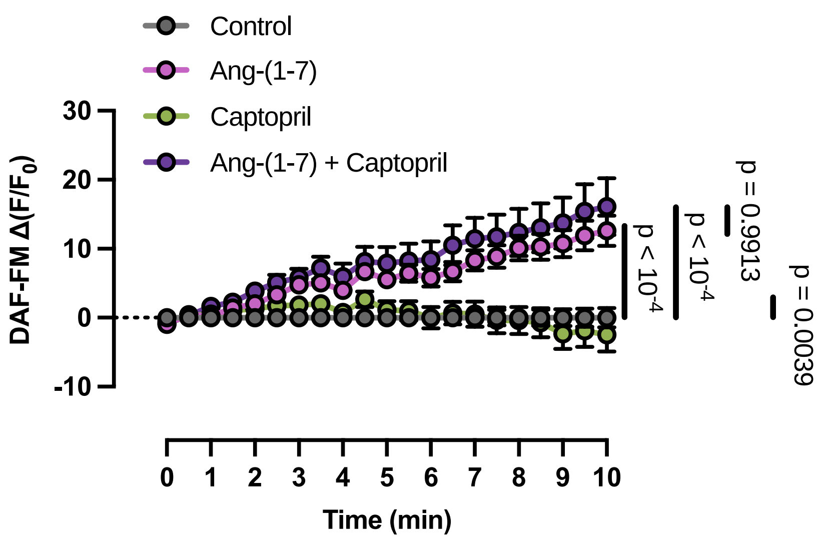


**Supplemental Figure S1 – Effects of Ang-(1-7) do not depend on its conversion to Ang-(1-5)**

Ang-(1-7) (100 nM) significantly increased NO release in HAEC when compared to vehicle treated cells. Prevention of conversion of Ang-(1-7) to Ang-(1-5) by ACE by addition of the ACE inhibitor captopril (100 nM) did not change the effect of Ang-(1-7) thus indicating that it is really Ang-(1-7) eliciting this effect and not its metabolite, Ang-(1-5). Data are presented as mean ± SEM from 3 independent experiments. Data were analysed by two-way RM-ANOVA. Differences were considered statistically significant when $p \leq 0.05$. Exact p-values are provided within the figure.

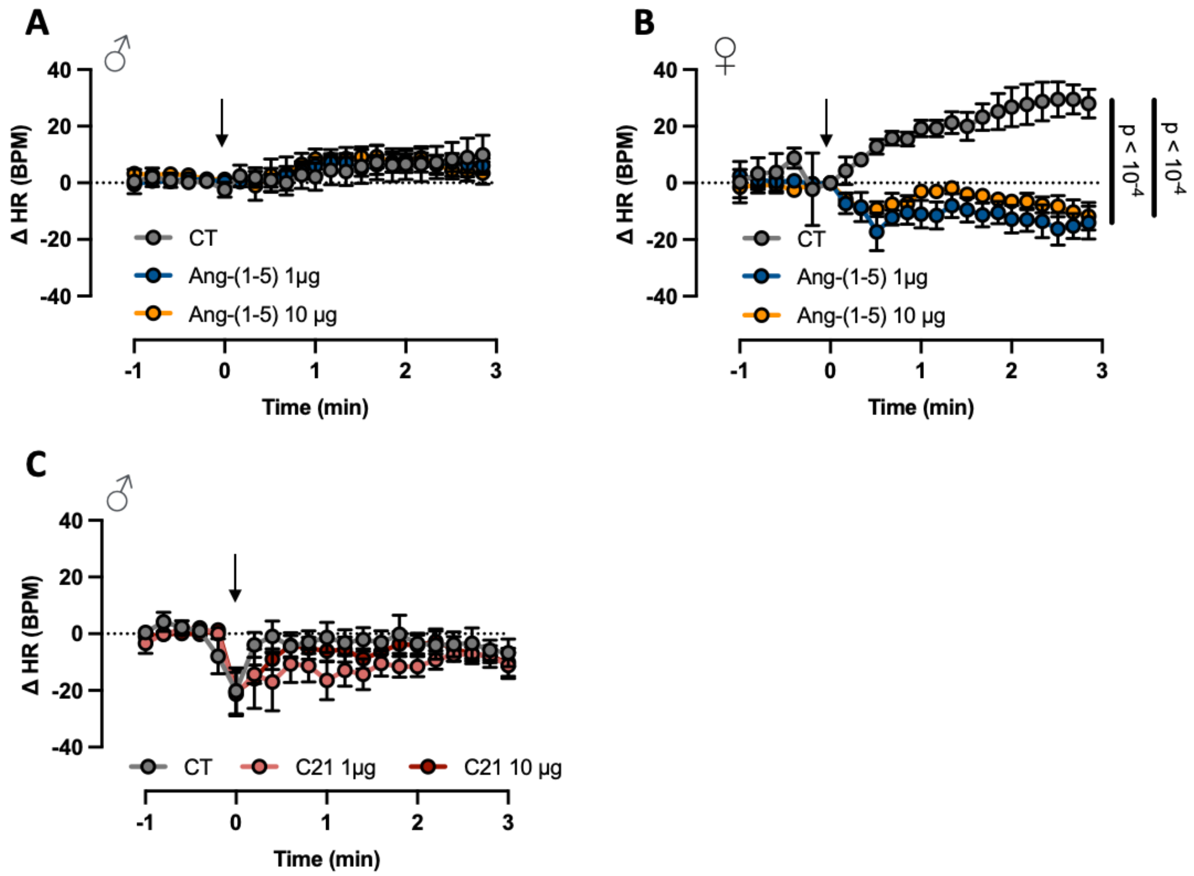


**Supplemental Figure S2 – Effects of Ang-(1-5) and C21 bolus injection on heart rate in anesthetized normotensive mice**

**A:** A bolus injection of Ang-(1-5) (1 or 10µg iv) had no effect on heart rate in anesthetized, normotensive male C57BL/6 mice. **B:** In female C57BL/6 mice, there was a statistically significant increase in HR after injection of vehicle (saline) in the control animals, which did not occur in animals treated with Ang-(1-5) (1 or 10µg iv). **C:** C21 (1 or 10µg iv) did not modify HR in anesthetized, normotensive male C57BL/6.

All data are shown as mean ± SEM and are from 9 mice/group (males, panel **A**), 4 mice/group (females, panel **B**) and 5 mice/group (males, panel **C**). Data were analysed by two-way repeated measure (RM)-ANOVA. Differences were considered statistically significant when $p \leq 0.05$. Exact p-values are provided within the figure.

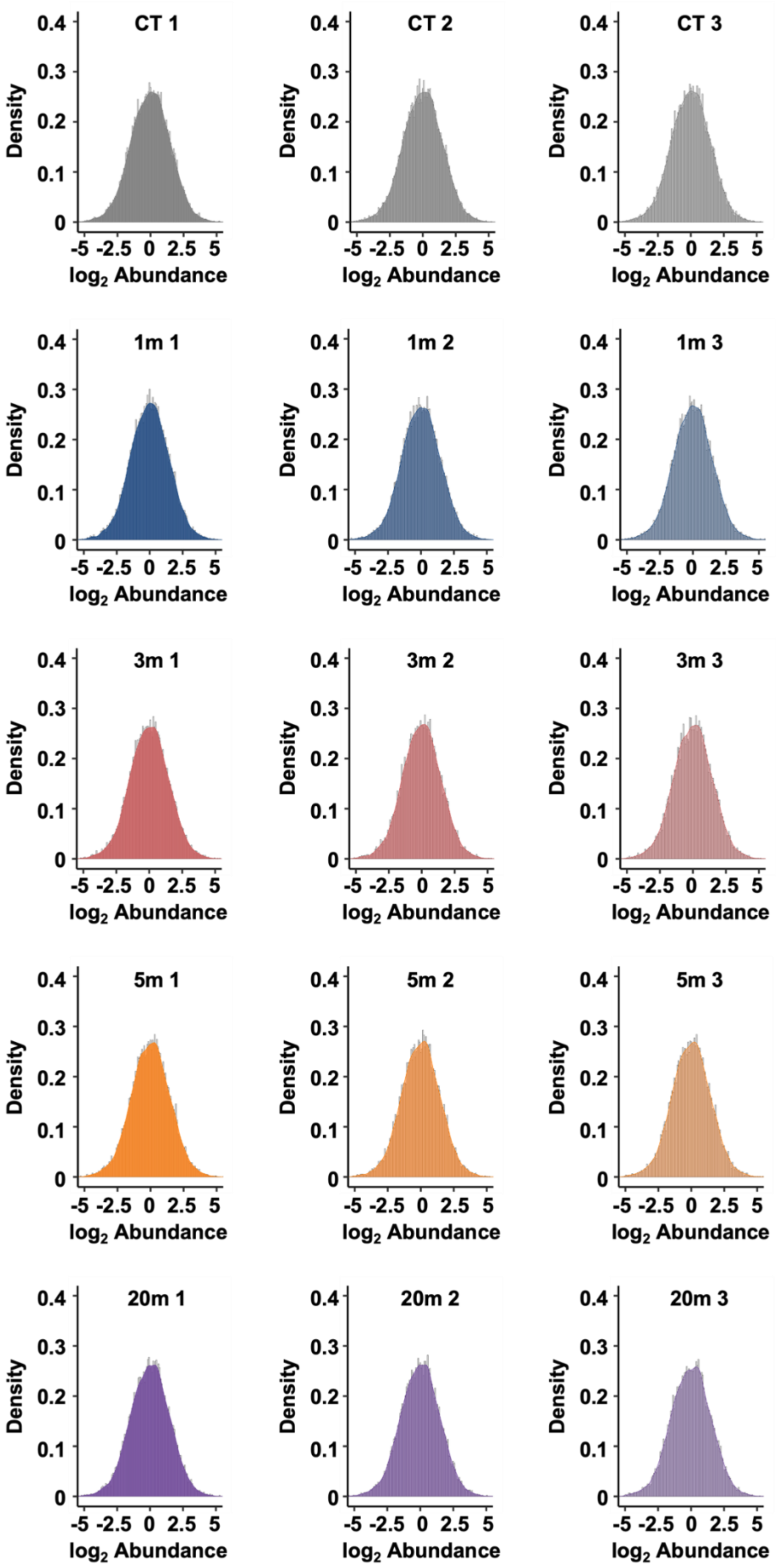
CT 1
CT 2
CT 3
1m 1
1m 2
1m 3
3m 1
3m 2
3m 3
5m 1
5m 2
5m 3
20m 1
20m 2
20m 3
Density
0.4
0.3
0.2
0.1
0
-5 -2.5 0 2.5 5
log2 Abundance

**Supplemental Figure S3 – Distribution of normalized phosphopeptide abundances within the phosphoproteome dataset**

Phosphopeptide abundances (shown as normalized $log_2$ values) in HAEC treated with Ang-(1-5) or vehicle (CT) for 1, 3, 5 or 20 minutes (3 biological replicates per timepoint) represented as histograms. The x-axis represents the log2-transformed abundances, the y-axis represents the density of the $log_2$-transformed phosphopeptide abundances. The abundances of all replicates from the phosphoproteome dataset follow a normal distribution.

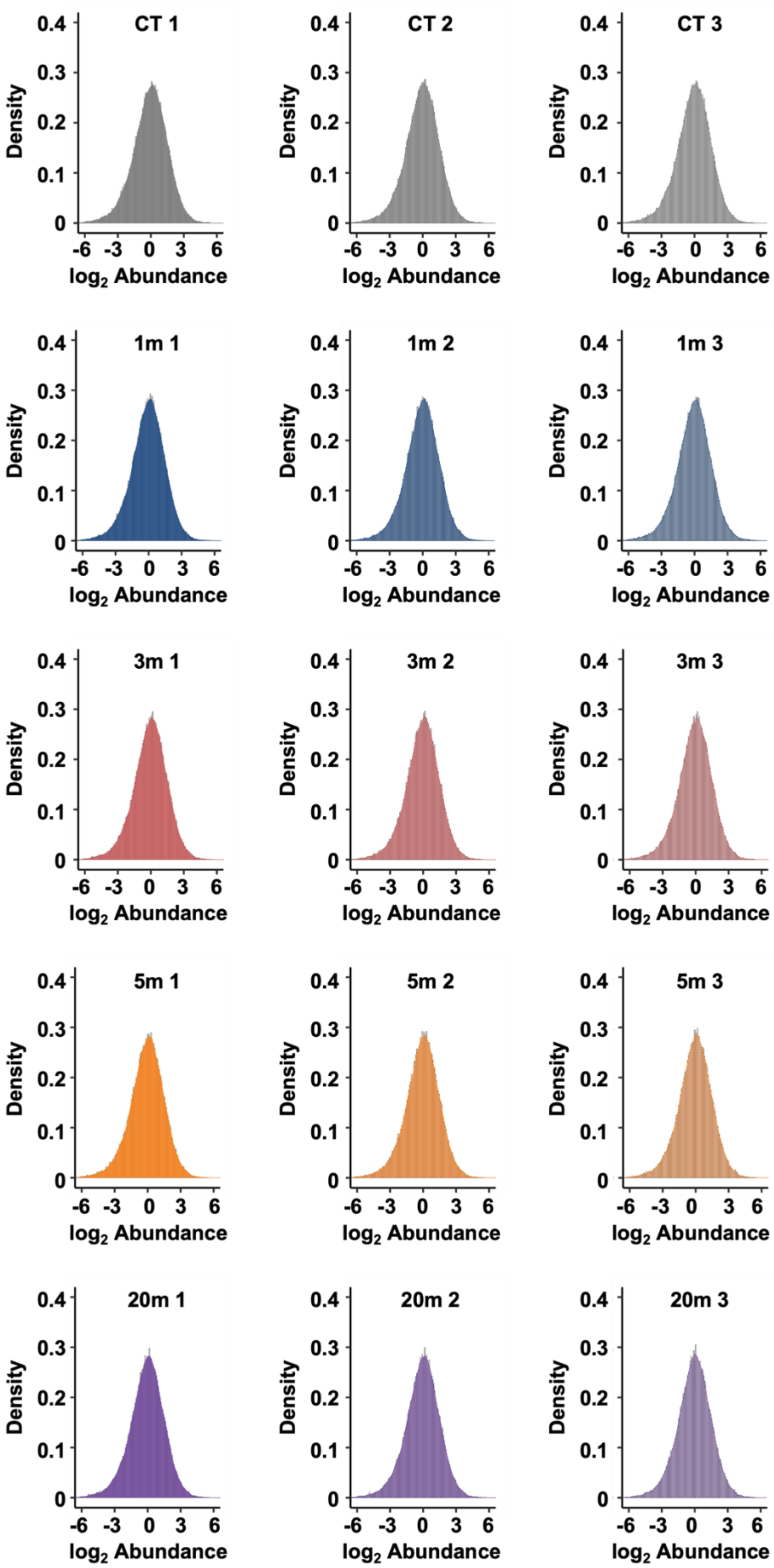
CT 1
CT 2
CT 3
1m 1
1m 2
1m 3
3m 1
3m 2
3m 3
5m 1
5m 2
5m 3
20m 1
20m 2
20m 3
Density
0.4
0.3
0.2
0.1
0
-6
-3
0
3
6
log2 Abundance

**Supplemental Figure S4 – Distribution of normalized non-modified peptide abundances within the proteome dataset**

Non-modified peptide abundances (shown as normalized $log_2$ values) in HAEC treated with Ang-(1-5) or vehicle, (CT) for 1, 3, 5 or 20 minutes (3 biological replicates per timepoint) represented as histograms. The x-axis represents the log2-transformed abundances, the y-axis represents the density of the $log_2$-transformed non-modified peptide abundances. The abundances of all replicates from the proteome dataset follow a normal distribution.
